# Effector dsRNA delivery via MgFe-layered double hydroxide nanocarriers confers prolonged protection against powdery mildew in pea

**DOI:** 10.64898/2026.01.02.697344

**Authors:** Poonam Ray, Mehak Bansal, Sumit Sagar, Bonamali Pal, Divya Chandran

## Abstract

Layered double hydroxides (LDHs) have emerged as promising nanocarriers for RNA interference (RNAi)-based crop protection due to their ability to stabilize and effectively deliver pathogen-specific double-stranded RNA (dsRNA). However, their potential against obligate biotrophic pathogens like powdery mildews remains unexplored. In this study, we investigated Magnesium Iron-LDH (MgFe-LDH) nanomaterials as carriers of fungal effector dsRNA (Ef-dsRNA) and evaluated the efficacy of the resulting complex in suppressing *Erysiphe pisi* (powdery mildew) infection in pea (*Pisum sativum*) via foliar spray application. The synthesized LDH nanomaterial exhibits strong leaf adherence, biocompatibility, and high dsRNA loading capacity, protects the dsRNA against RNase-mediated degradation, and facilitates its sustained release over time. Foliar spray application of the Ef-dsRNA-LDH complex on intact pea plants results in enhanced gene silencing of the target fungal effector and confers greater and prolonged local and systemic powdery mildew disease protection compared to treatments with dsRNA or LDH alone. Following spray application, the LDH nanomaterial is rapidly taken up by intact pea leaf cells and from the plant into *E. pisi* hyphae via haustoria, enabling efficient dsRNA delivery and silencing of the target gene up to 15 days post-application. Overall, our Ef-dsRNA-LDH formulation offers a robust, sustainable, and targeted approach for RNAi-mediated control of powdery mildews.

## Introduction

Phytopathogenic fungi represent a significant threat to global agriculture due to their epidemic potential, driven by high virulence, broad host ranges, and adaptability to adverse environmental conditions, coupled with the capacity to evolve new lineages (Fisher *et al*., 2012; Stukenbrock & Gurr, 2023). Fungal outbreaks disproportionately affect food security in underdeveloped regions, where restrictive trade policies can exacerbate crop price volatility (Godfray *et al*., 2016). Pea (*Pisum sativum*) is a critical food crop and a significant source of protein in vegetarian diets (Lefranc-Millot & Teichman-Dubois, 2019). Powdery mildew (PM), primarily caused by the biotrophic fungal pathogen *Erysiphe pisi*, poses a major threat to pea production, with yield losses ranging from 25–70% (Jha *et al*., 2019; Maria *et al*., 2024).

Management of PM involves the use of chemical and biological fungicides and genetically resistant crop varieties (Fondevilla & Rubiales, 2012). While chemical fungicides are effective, their utility is influenced by environmental factors and application protocols, and they often cause adverse effects on non-target organisms, human health, and plant physiology (Wan *et al*., 2025; Pearson *et al*., 2016). Additionally, fungicide overuse often promotes the emergence of resistant fungal strains (Vielba-Fernandez *et al*., 2020). Biocontrol agents offer an eco-friendly alternative, though their efficacy heavily depends on environmental conditions (Ons *et al*., 2020). To date, the use of genetically resistant plants remains the most sustainable approach; however, the limited pool of PM resistance genes in pea and the non-acceptance of GMOs by the general public pose challenges for developing resistant plants (Brown, 2015; Devi *et al*., 2022; Bekele-Alemu *et al*., 2025).

RNA interference (RNAi)-based approaches, including host-induced (HIGS) and spray-induced (SIGS) gene silencing, have emerged as environment-friendly alternatives to traditional fungicides for managing fungal diseases (Ray *et al*., 2022; Mann *et al*., 2023). In these technologies, double-stranded (ds) RNA targeting fungal genes is applied to plants as a pre-treatment, creating a protective shield for a specific period, allowing for siRNA production and suppression of infection-related fungal genes. These strategies target essential fungal genes to reduce fungal growth (Gebremichael *et al*., 2021; Ray *et al*., 2022). SIGS offers an advantage over HIGS by circumventing the need for crop-specific transformation protocols and providing greater flexibility, as it is not confined to specific gene targets or applications (Taning *et al*., 2020).

Previously, we demonstrated that silencing an *E. pisi ribonuclease-like* effector via dsRNA infiltration in pea leaves disrupts the fungal infection process (Sharma *et al*., 2019). Targeting fungal effectors through SIGS offers a pathogen-specific strategy for disease management, as these genes are crucial for suppressing host immunity but are generally non-essential for fungal viability *in vitro* (Ray *et al*., 2022; Ouyang *et al*., 2025). In contrast, structural genes like tubulin are highly conserved and often present in multiple isoforms, making them less effective due to functional redundancy and more prone to off-target risks. A key limitation of SIGS is the transient nature of protection conferred by dsRNA, owing to its susceptibility to RNase degradation, absence of sustained release mechanisms, limited uptake of dsRNA by plant and pathogen, and loss due to environmental factors such as rainfall (Secic & Kogel, 2021; Venu *et al*., 2024). Therefore, controlled delivery systems are required to enhance the stability and ensure controlled release of dsRNA to prolong the RNAi-mediated protection (Ray *et al*., 2022; Quilez-Molina *et al*., 2024).

Layered double hydroxides (LDH) offer a promising solution for delivering dsRNA. These biodegradable nanocarriers, consisting of divalent (e.g., Mg, Zn) and trivalent (e.g., Fe, Al) metal ions, feature positively charged hydroxide layers intercalated with anions (Cl^-^, NO_3-_, etc.), enabling efficient delivery of negatively charged biomolecules such as dsRNA (Zhang *et al*., 2021; Singha Roy *et al*., 2022). Studies have demonstrated the utility of Magnesium Aluminium-LDH (MgAl-LDH) in dsRNA delivery, providing sustained protection against viruses (Mitter *et al*., 2017). However, concerns regarding aluminium toxicity in plants and soil (Chandran *et al*., 2008; Bojorquez-Quintal *et al*., 2017) led to the development of MgFe-LDH, which has demonstrated efficacy in protecting crops against pests such as *Bemisia tabaci* (white fly) (Jain *et al*., 2022) and necrotrophic fungal pathogens such as *Botrytis cinerea* (Niño-Sánchez *et al*., 2022) and *Sclerotinia sclerotiorum* (Mukherjee *et al*., 2024).

In this study, we demonstrate the efficacy of MgFe-LDH as a delivery system for *E. pisi* effector-targeting double-stranded RNA (Ef-dsRNA) for RNAi-based protection against PM disease in pea. We prove that hexagonally structured, layered MgFe-LDH nanomaterials possess a net positive charge, enabling efficient loading of dsRNA. The LDH protects dsRNA from RNase-mediated degradation and facilitates sustained release of *E. pisi* Ef-dsRNA under ambient conditions. The LDH nanomaterial gradually degrades into Mg²⁺ and Fe³⁺ ions on the pea leaf surface and exhibits biocompatibility across multiple plant species. Additionally, we show that the MgFe-LDH nanomaterial is efficiently internalized by pea and other plants, and is, importantly, also taken up by *E. pisi* hyphae from the leaf, likely through haustoria. The spray application of partially-loaded Ef-dsRNA-LDH complexes reveals dual-phase protection, an immediate antifungal response driven by free dsRNA and prolonged protection mediated by the gradual release of dsRNA from the complex. The enhanced stability imparted by LDH further facilitates the systemic movement of dsRNA into newly emerging leaves, contributing to systemic resistance. Our findings position MgFe-LDH as a robust, biodegradable platform for RNAi-based PM control in pea and other crops.

## Results

### Synthesized MgFe-LDH forms hexagonal nanosheets

Transmission electron microscopy (TEM) revealed well-defined hexagonal platelets of MgFe-LDH with lateral sizes ranging from 60–150 nm (Fig. **S1a**). High-resolution TEM (HRTEM) displayed clear lattice fringes, one corresponding to the (009) plane with an interplanar spacing of 0.282 nm (Fig. **S1b**). Field emission scanning electron microscopy (FE-SEM) further confirmed the hexagonal morphology and showed both horizontal and vertical orientations of the stacked platelets, characteristic of their lamellar structure (Fig. **S1c–d**) (Silva Neto et al., 2021). Energy-dispersive X-ray spectroscopy (EDS) mapping demonstrated a uniform distribution of magnesium (Mg), iron (Fe), and oxygen (O) across the nanomaterial surface (Fig. **S1e**).

X-ray photoelectron spectroscopy (XPS) confirmed the presence of Mg, Fe, O, Cl, and C (Fig. **S1f**). The Mg 1s peak at 1303.66 eV indicated divalent Mg²⁺ (Fig. **S1g**), while Fe 2p peaks at 712.49 eV and 726.42 eV, with a satellite at 724 eV, confirmed Fe³⁺ (Fig. **S1h**). O 1s peaks at 531.26 eV and 532.45 eV corresponded to metal hydroxides and hydrogen-bonded water, respectively (Fig. **S1i**). Cl 2p peaks were also resolved into two individual peaks of Cl 2p1/2 at 199.56 eV and Cl 2p3/2 at 197.35 eV (Fig. **S1j**). FTIR analysis showed characteristic bands for -OH groups, intercalated water, and NO₃⁻ vibrations at 3400, 1640, and 1380 cm⁻¹, respectively (Fig. **S1k**) (Hidayati et al., 2019). XRD analysis showed sharp reflections at 11.29°, 22.42°, 34.30°, 59.42°, and 60.98°, indexed to (003), (006), (009), (110), and (113) planes, confirming layered structure and anion intercalation (Fig. **S1l**). BET analysis showed a surface area of 21.96 m²/g, pore diameter of 3.825 nm, and pore volume of 0.0367 cm³/g (Fig. **S1m**).

### MgFe-LDH loads, protects, and releases dsRNA via degradation

To evaluate the dsRNA loading efficiency of the synthesized MgFe-LDH, various dsRNA-LDH mass ratios were analyzed via agarose gel electrophoresis. Successful loading, indicated by the retention of dsRNA in the well, was observed with increasing LDH concentration, and complete loading was observed at a 1:50 dsRNA:LDH mass ratio (Fig. **1a**). Residual unbound dsRNA was quantified via ImageJ, using 1 µg of naked dsRNA (relative intensity = 1.0) as a reference. The relative intensities decreased from 0.98 (1:1) to 0.00 (1:50), confirming efficient dsRNA loading by MgFe-LDH in a mass-dependent manner. Additionally, after dsRNA loading, the hydrodynamic size of the LDH increased from an average of 337 ± 9 nm to 463 ± 41, whereas the zeta potential decreased from 25 ± 1.5 mV to 15 ± 0.8 mV (Fig. **1b**). The polydispersity index (PDI) increased from 0.23 to 0.49 after dsRNA loading (Fig. **1b**), likely due to the size variation between the dsRNA-loaded and unloaded LDH. TEM analysis revealed thin, fine projections on the LDH surface, along with an increase in the size of the nanomaterial from an average of 80 ± 20 nm to 180 ± 45 nm (Fig. **S2)**.

**Fig. 1.**
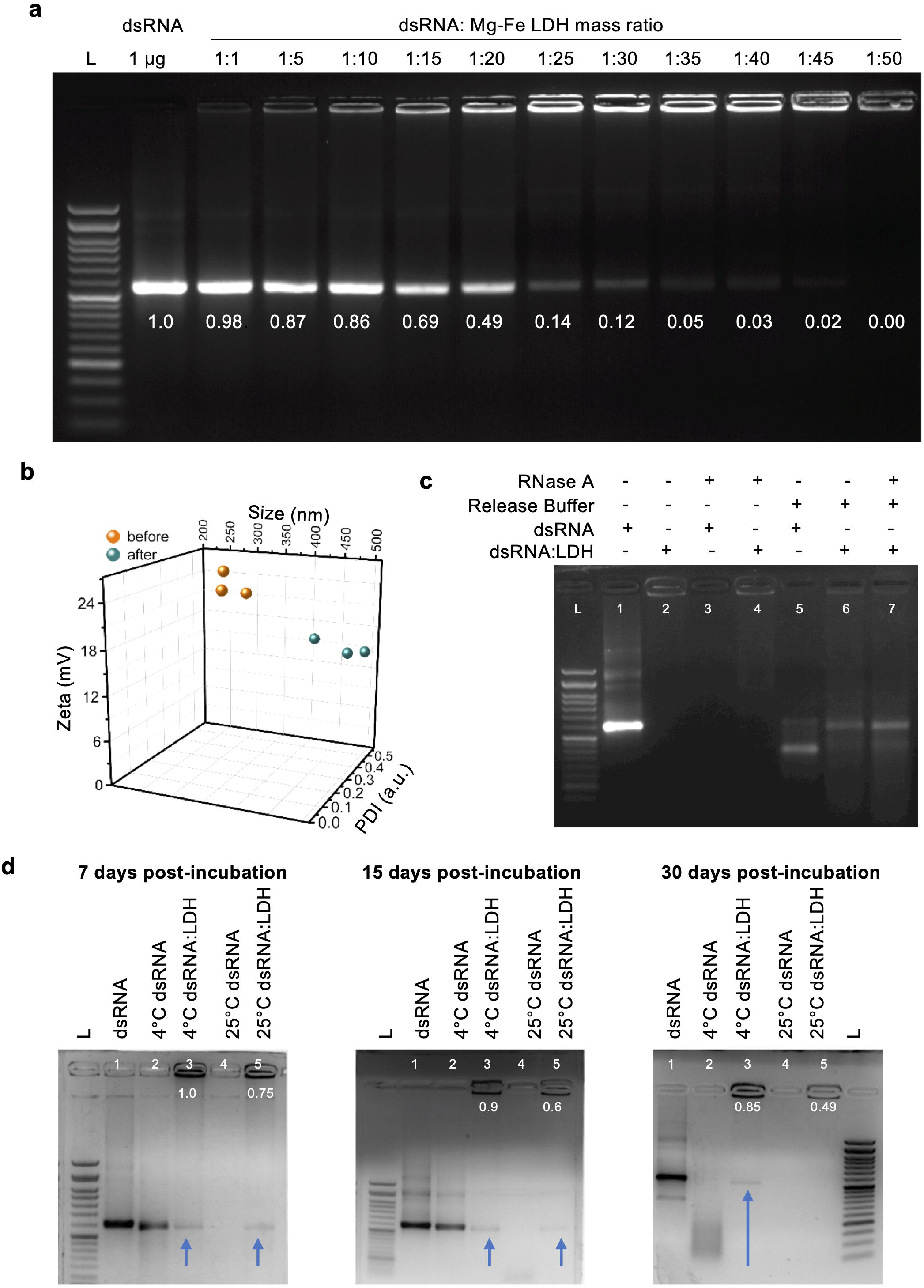
dsRNA loading capacity of MgFe-LDH and release kinetics. (a) Loading of dsRNA at different dsRNA-LDH mass ratios; L=50 bp ladder. Values shown below the dsRNA bands in the gel indicate the ratio of band intensity in each lane to the band intensity of dsRNA alone (taken as 1) (b) DLS measurement of LDH size, PDI, and zeta potential before and after dsRNA loading (c) Treatment of dsRNA and dsRNA:LDH (1:50) with RNase A. The dsRNA from treated and untreated dsRNA-LDH samples was released using release buffer (pH-3) before gel electrophoresis; L=50 bp ladder (d) Temperature-dependent release of dsRNA from dsRNA-LDH (1:50) samples at different time points after incubation; L=50 bp ladder. Blue arrows indicate the released dsRNA.

To test whether LDH protects the dsRNA, naked dsRNA and fully loaded dsRNA:LDH (1:50) complex were treated with RNase A, subsequently released, and analyzed via agarose gel electrophoresis. Naked dsRNA was fully degraded following RNase A treatment (Fig. **1c**, lane 3), whereas LDH-bound dsRNA remained intact (lane 7). The addition of the release buffer converted a significant proportion of the naked dsRNA, but not the LDH-bound dsRNA, into single-stranded RNA (ssRNA) (lane 5). These results suggest that the LDH protects the dsRNA from RNase activity and preserves its structural integrity at low pH.

Next, we evaluated the temperature-dependent degradation of LDH and dsRNA release at two temperatures (4°C and 25°C) and three time points (7, 15, and 30 days). These temperatures were selected keeping in view the minimum temperature at which pea seeds can germinate (4°C) and the optimal temperature for pea growth (22-25°C) (Olivier & Annandale, 1998). Although dsRNA release occurred at both temperatures, LDH degraded faster at 25°C, as indicated by reduced band intensity in the gel (Fig. **1d**). Further, the released dsRNA was more stable at 4°C, as the intensity of the dsRNA band in the gel gradually diminished with time at 25°C but not at 4°C (Fig. **1d**). These results suggest that the LDH degrades faster and possibly releases a larger fraction of the bound dsRNA at higher temperatures, whereas the complex remains stable for longer at lower temperatures.

### Spray application of Effector-dsRNA-loaded LDH offers greater and prolonged protection against PM

We previously identified a ribonuclease-like gene (*Ep*CSEP001) as the highest expressed *E. pisi* effector candidate and demonstrated that dsRNA infiltration-mediated silencing of this gene delayed PM disease progression on pea leaves by 3-5 days (Sharma *et al*., 2019). Thus, we selected *EpCSEP001* (referred to as *Ef*) as the target fungal gene to evaluate the efficacy of MgFe-LDH as a dsRNA nanocarrier and its potential to extend the protection window against PM. A pilot study identified 200 ng/μL as the optimal *Ef*-dsRNA dose for crop protection assays, as spray application of this concentration reduced fungal hyphae number at 3 days post-inoculation (dpi) (Fig. **S3a,b**) and PM symptom development at 7 dpi compared to the GFP-dsRNA control treatment (Fig. **S3c,d**). Accordingly, a significant reduction in target *Ef* gene expression and fungal load was observed until 5 dpi (Fig. **S3e**). Additionally, no visible signs of phytotoxicity were observed at 200 ng/μL, whereas a higher dose (500 ng/ μL) occasionally caused leaf wilting (Fig. **S3d**).

To assess the effectiveness of the Ef-dsRNA-LDH formulation in suppressing PM growth on pea, we used a dsRNA:LDH loading mass ratio of 1:20, in which ∼50% dsRNA is loaded on the LDH, with the expectation that the unbound portion would offer immediate protection upon spraying. Pea leaves were sprayed with GFP-dsRNA, Ef-dsRNA, LDH, GFP-dsRNA-LDH, or Ef-dsRNA-LDH and inoculated with *E. pisi* after 24 hours. Compared to the GFP and LDH treatments, pea leaves sprayed with Ef-dsRNA alone or the Ef-dsRNA-LDH complex exhibited reduced PM disease symptoms at all infection time points (7, 10, and 15 dpi), with the percent reduction always greater in the Ef-dsRNA-LDH-sprayed leaves (Fig. **2a,b**). Correspondingly, *Ef* expression was significantly down-regulated at all infection time points in the Ef-dsRNA-LDH-sprayed leaves (Fig. **2c**). Dot blot analysis revealed that Ef-dsRNA remains stable for extended periods when delivered with the LDH, with Ef-dsRNA detected up to 15 dpi in Ef-dsRNA-LDH-sprayed leaves compared to 7 dpi in Ef-dsRNA-sprayed leaves (Fig. **2d**). These findings indicate that the Ef-dsRNA-LDH complex affords greater protection against PM in pea than the Ef-dsRNA alone.

**Fig. 2.**
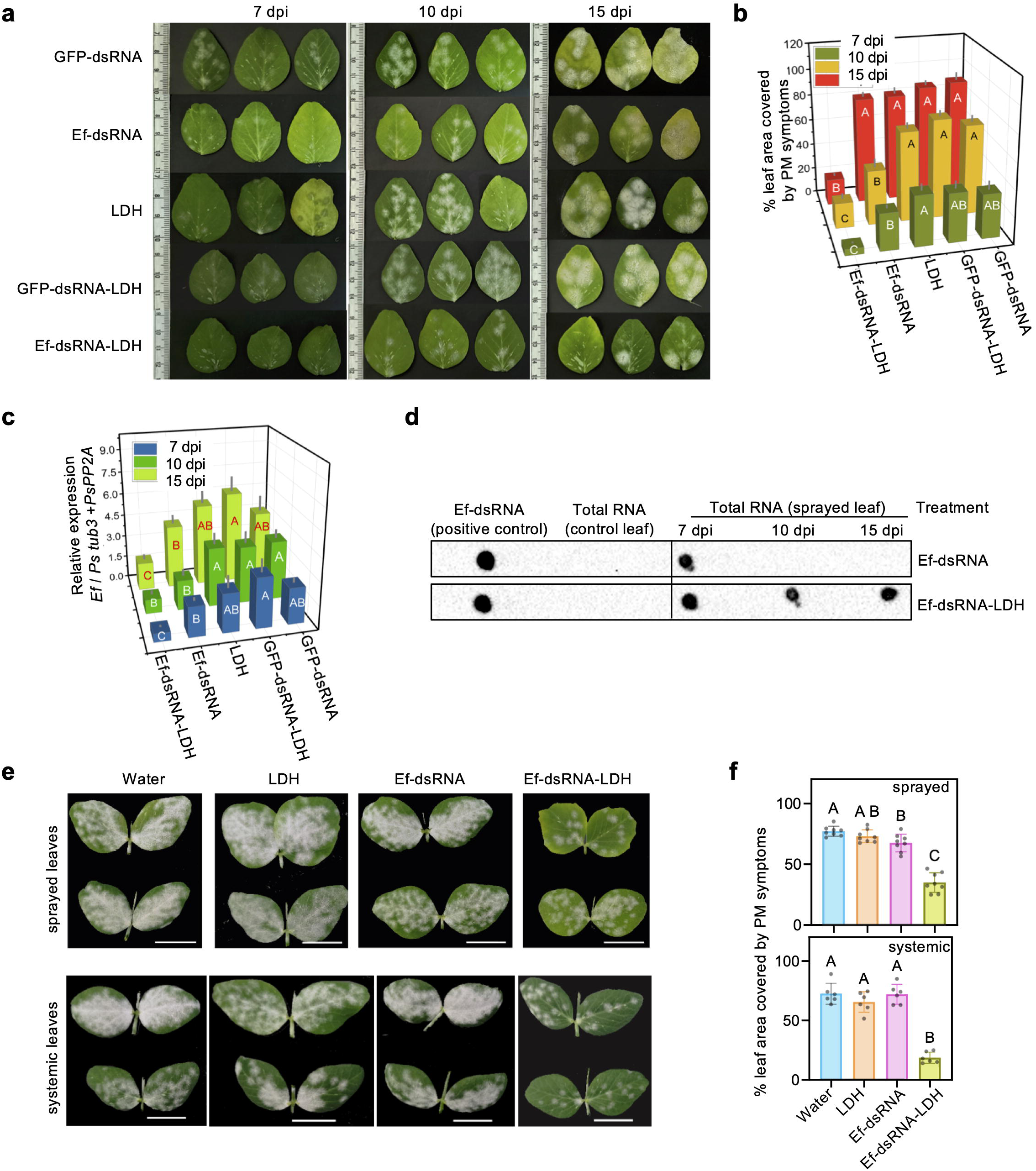
Spray treatment of Ef-dsRNA-LDH offers prolonged and systemic protection against PM in pea. (a) White powder-like disease symptoms on pea leaves at 7, 10, and 15 days post-inoculation (dpi) with *E. pisi*. Leaves of 15-day-old pea plants were sprayed with LDH, Ef/GFP-dsRNA, or Ef/GFP-dsRNA-LDH on day 0 (n = 6 leaves per treatment group) and inoculated with *E. pisi* one day post-treatment. (b) Percent (Mean ±SD) leaf area covered by PM disease symptoms quantified using ImageJ (c) Relative expression of the target *Ef* gene. Data represent mean (±SD) relative expression of *Ef* gene normalized to the endogenous controls *Pstubulin* and *PsPP2A*. (d) Representative dot blot images showing the presence of Ef-dsRNA in Ef-dsRNA or Ef-dsRNA-LDH-sprayed leaves at different time points after *E. pisi* inoculation. 20 ng of Ef-dsRNA was spotted as the positive control, and 2 µg total RNA from unsprayed leaves was spotted as the negative control. (e) PM disease symptoms on sprayed and unsprayed, systemic pea leaves at 10 days post-inoculation (dpi) with *E. pisi*. Leaves of 15-day-old pea plants were sprayed with water, LDH, Ef-dsRNA, or Ef-dsRNA-LDH (1:20) on day 0 (n = 6 leaves per treatment group) and inoculated with *E. pisi* 15 days post-spray; Scale = 2 cm. (f) Bar graphs showing percent leaf area (mean ±SD) covered by PM disease symptoms quantified from images in (e) using Image J. Statistical significance was computed via One-way ANOVA (*p* < 0.0001) along with Tukey’s multiple comparisons test denoted as uppercase letters. Three independent experiments gave similar results.

To determine whether Ef-dsRNA-LDH provides long-term protection against PM, pea plants were sprayed with Ef-dsRNA, LDH, Ef-dsRNA-LDH, or water, and challenged with *E. pisi* 15 days after application. Compared to the naked Ef-dsRNA, leaves treated with Ef-dsRNA-LDH exhibited significantly reduced PM disease symptoms on both sprayed and newly emerged unsprayed leaves (Fig. **2e,f**), indicating that Ef-dsRNA-LDH provides prolonged and systemic protection against PM in pea.

### LDH adheres to the leaf surface and internalizes into pea cells and fungal hyphae

We assessed whether the extended protective effect of the Ef-dsRNA-LDH correlates with better adherence and internalization of the complex into pea leaf tissues. For this, MgFe-LDH was labelled with FITC, and successful labelling was confirmed through DLS, FTIR, and TEM. DLS analysis indicated an increase in hydrodynamic size and a decrease in zeta potential of the FITC-LDH relative to the unlabelled LDH (Fig. **S4a**). FTIR analysis corroborated the labelling, revealing increased spectral waviness in the range of 1454–2002 cm⁻¹ and reduced peak intensity at the wavenumber 2981 cm⁻¹, indicative of changes in functional groups after successful labelling (Fig. **S4b**). TEM analysis showed significant differences in surface contrast between FITC-LDH and unlabelled LDH (Fig. **S4c**).

To assess the adherence of LDH on the leaf surface, pea leaves sprayed with water, FITC, or FITC-LDH were incubated in the dark for 1 hour, washed thoroughly with water, and imaged under a handheld UV lamp at 1 and 12 hours post-wash. A stronger green FITC fluorescence signal was observed at both time points in the FITC-LDH-sprayed leaves compared to those sprayed with FITC alone (Fig. **3a**), indicating robust adherence of LDH on the leaf surface.

**Fig. 3.**
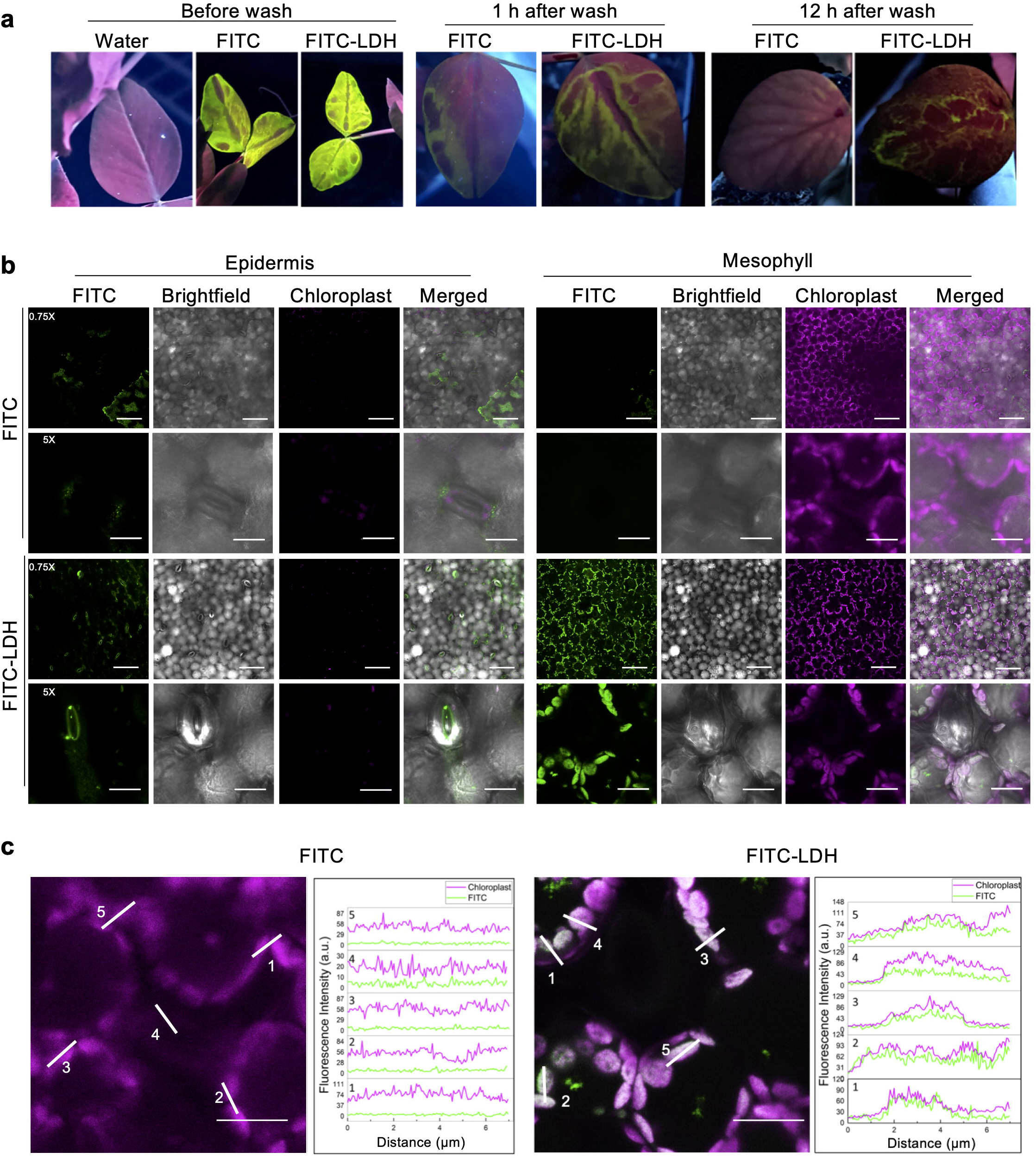
Adherence and internalization of MgFe-LDH in pea leaves after spray application. (a) Pea leaves were sprayed with FITC-labelled LDH (1:20) or FITC alone and rinsed vigorously with water. Leaves were visualized under a handheld UV lamp before wash (left panel) and 1 h and 12 h post-wash (right panel). FITC fluorescence is shown in green. (b) Representative confocal images of LDH internalization into epidermal and/or mesophyll cells of pea leaves. The adaxial surface of pea leaves was sprayed with FITC-LDH or FITC alone, rinsed with water after 1 h, and observed immediately under a 40x oil objective of a confocal microscope at magnification 0.75x, Scale = 80 μm, and magnification 5x, Scale = 15 μm. FITC fluorescence is shown in green and chloroplast autofluorescence in magenta. (c) Zoomed-in view of the mesophyll cells of FITC and FITC-LDH sprayed pea leaves, along with line scans (white lines 1-5) showing an overlap in fluorescence intensity of FITC (green) and chloroplast (magenta) signals in FITC-LDH sprayed samples; Scale = 10 μm

To assess the internalization of the LDH into pea leaf cells, FITC-LDH and FITC-sprayed pea leaves were analysed via confocal microscopy 1 hour post-wash. Green fluorescence from FITC, magenta chloroplast autofluorescence (serving as a marker for mesophyll cell localization), and brightfield images were merged to demonstrate LDH uptake in leaf epidermal and mesophyll layers. Compared to the FITC-sprayed leaves, a stronger FITC fluorescence signal was visible in the epidermal layer of FITC-LDH-sprayed leaves, especially around the stomata (Fig. **3b**, left panel). No detectable fluorescence signal was observed within the mesophyll region of leaves sprayed with FITC alone. In contrast, leaves sprayed with FITC-LDH exhibited green fluorescence within the mesophyll layer, with nearly all cells displaying fluorescence (Fig. **3b**, right panel), suggesting effective penetration of FITC-LDH in the targeted tissues. To further confirm the uptake of FITC-LDH into mesophyll cells, line profile analysis was performed across regions showing colocalization between the FITC signal and chloroplast autofluorescence, and the corresponding fluorescence intensity values were plotted. The overlapping fluorescence profiles of chloroplasts and FITC-LDH (Fig. **3c**, right panel) in the FITC-LDH-sprayed pea leaves, but not in FITC-sprayed leaves (Fig. **3c**, left panel), indicate successful internalization of the FITC-LDH into mesophyll cells. Similar experiments conducted in *Nicotiana benthamiana* (Fig. **S5**) and rice (Fig. **S6**) yielded comparable results, validating the consistency and reproducibility of FITC-LDH uptake across different plant species.

To determine whether the MgFe-LDH is also taken up by fungal hyphae, leaves of pea plants were sprayed with FITC alone or FITC-LDH and subsequently inoculated with *E. pisi*. Confocal imaging was performed at 12, 24, 48, and 72 hpi to visualize FITC-LDH signal (green) within fungal hyphae stained with Calcofluor White (blue). At the early infection stages (12 and 24 hpi), the green FITC fluorescence signal was visible within the fungal conidia and emerging hypha in both FITC and FITC-LDH-sprayed leaves (Fig. **4a**). At later infection stages (48 and 72 hpi), the green FITC fluorescence signal appeared strongly localized at fungal haustorial sites (marked with white arrows) and within secondary hyphae in the FITC-LDH-sprayed leaves than in leaves sprayed with FITC alone (Fig. **4a**). A 3D surface reconstruction of the fungal hyphae with solid fill or 60% transparency was performed using the 72 hpi FITC-LDH images to demonstrate internalization of LDH into fungal hyphae (Fig. **4b**). The FITC-LDH green fluorescence signal was only visible when the fungal surface was set to 60% transparency, implying that the signal is not on the surface but originates from inside the fungal hyphae. Further, the intense green fluorescence at focal points below the hyphae (Fig. **4a,b**) suggests active siphoning of FITC-LDH by the fungus from the pea leaf through haustoria formed within infected epidermal cells.

**Fig. 4.**
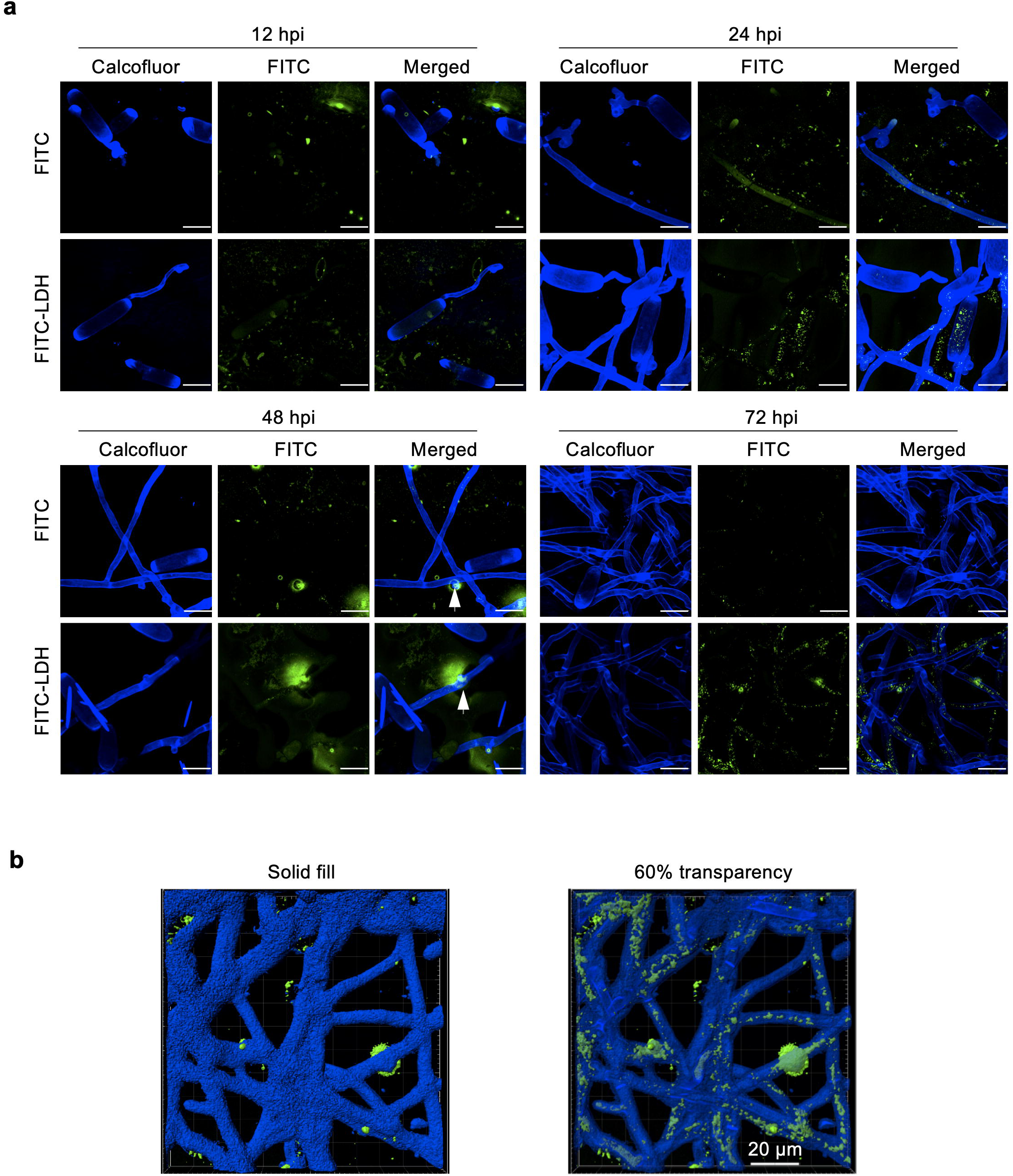
MgFe-LDH uptake by PM hyphae from pea leaves after spray application. (a) Representative confocal images showing uptake of FITC-LDH (1:20) or FITC alone into fungal hyphae at different hours post-inoculation (hpi) with *E. pisi* fungus. The adaxial surface of pea leaves was sprayed with FITC-LDH or FITC alone, rinsed with water after 1 h, and inoculated with *E. pisi* conidia. The fungus is stained with calcofluor (blue), and the FITC fluorescence is shown in green; Scale = 20 μm (b) Imaris-based 3D reconstruction of Z-stacks from 72 hpi image of FITC-LDH using solid fill colour (blue) or 60% transparency to show internalization of FITC-LDH (green) into fungal hyphae.

### LDH facilitates Ef-dsRNA uptake into the pea leaves and fungal hyphae

To ascertain whether MgFe-LDH facilitates dsRNA uptake into plant cells, leaves of pea plants were sprayed with Cy3-labelled Ef-dsRNA (dsRNA-Cy3), dsRNA-Cy3-LDH, or Cy3 alone, incubated in the dark for 1 hour, washed thoroughly with water, and returned to the dark for an additional 11 hours. The additional dark incubation period was designed to allow sufficient time for internalization of dsRNA-Cy3 and dsRNA-Cy3-LDH. The LDH was not labelled in these experiments because the surface charge (Fig. **S4a**) and, consequently, the dsRNA loading capacity of the nanomaterial substantially reduced upon FITC-labelling. Red fluorescence from Cy3, green chloroplast autofluorescence (serving as a marker for mesophyll cells), and brightfield images were merged to demonstrate dsRNA uptake in leaf epidermal and mesophyll layers (Fig. **5a**).

**Fig. 5.**
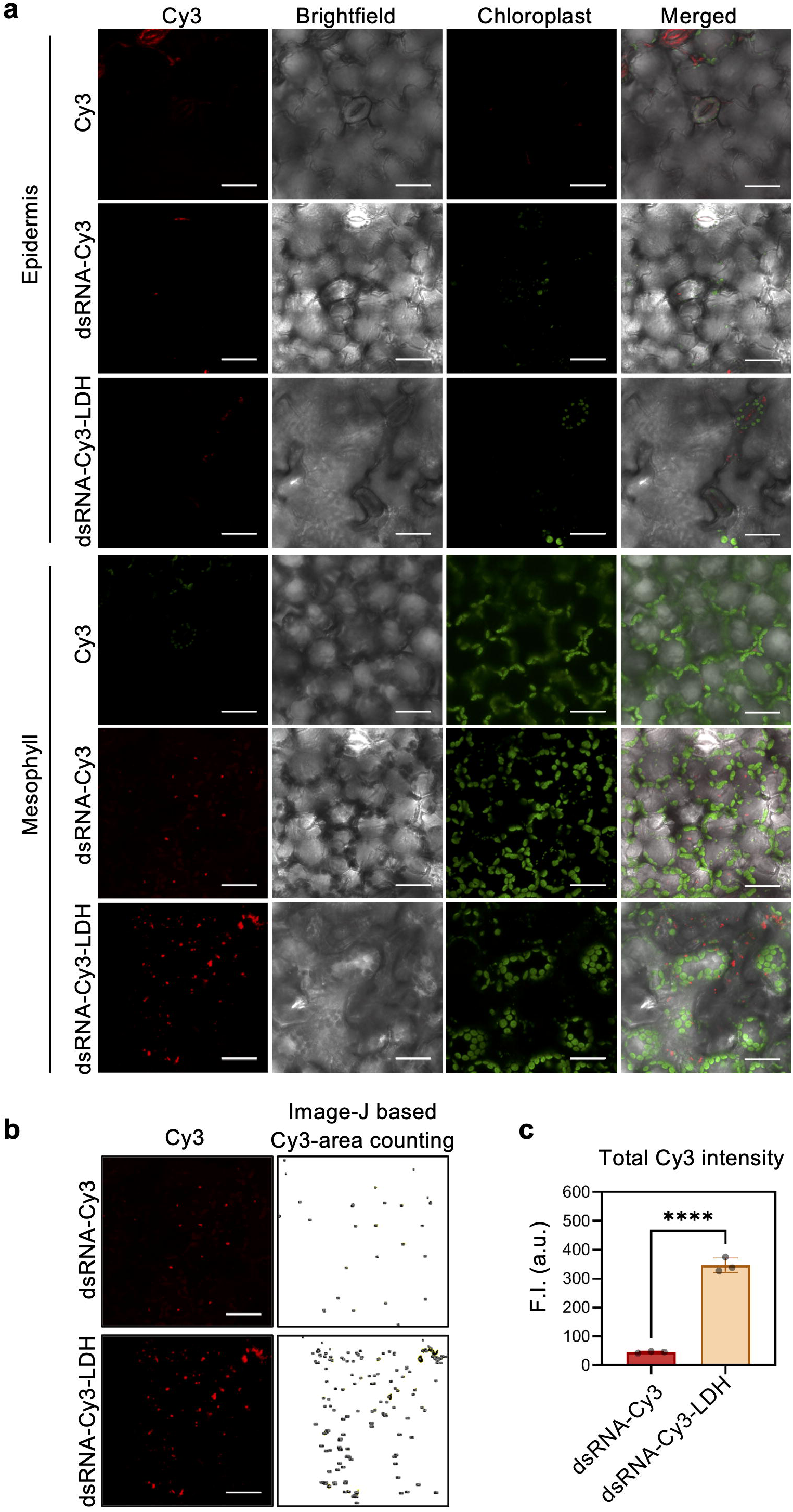
Internalization of dsRNA loaded on MgFe-LDH into pea leaves. (a) Representative confocal images showing dsRNA-Cy3 uptake into pea leaves at 12 hours post-spray. The adaxial surface of pea leaves was sprayed with Cy3 alone, dsRNA-Cy3, or dsRNA-Cy3-LDH (1:20), and the upper epidermal and mesophyll cells were imaged. Cy3 fluorescence is shown in red and chloroplast autofluorescence in green; Scale *=* 30 μm (b) Cy3 fluorescence intensity was measured using the ImageJ threshold setting from the mesophyll cells of dsRNA-Cy3 and dsRNA-Cy3-LDH sprayed pea leaves; Scale = 30 μm (c) Graphical representation of Cy3 fluorescence intensity as measured in ImageJ from three images. Three independent experiments were performed. Statistical significance was computed using an unpaired t-test (α < 0.05); **** = P ≤ 0.0001

In all three treatments, a faint Cy3 (red) fluorescence signal was detected in the leaf epidermal layer (Fig. **5a**). Comparatively, in the mesophyll layers, marked differences in the Cy3 signal intensity were visible between the three treatments, with the highest signal observed in the dsRNA-Cy3-LDH-sprayed leaves, followed by the leaves sprayed with dsRNA-Cy3 and Cy3 alone (Fig. **5a**). Quantification of the Cy3 fluorescence intensity revealed an ∼8-fold higher Cy3 signal in the dsRNA-Cy3-LDH-sprayed leaves than in the leaves sprayed with dsRNA-Cy3 (Fig. **5b,c**), suggesting that LDH enhances the uptake of the dsRNA into plant cells.

To assess the internalization of the dsRNA into fungal hyphae, leaves of pea plants were sprayed with dsRNA-Cy3, dsRNA-Cy3-LDH, LDH+Cy3, or Cy3 alone and inoculated with *E. pisi*. Confocal imaging was performed at 12, 24, 48, and 72 hpi to visualize dsRNA-Cy3 signal (red) within fungal hyphae stained with Calcofluor White (blue). At 12 and 24 hpi, Cy3 fluorescence was visible within fungal conidia or beneath the fungal appressorium in all treatments (Fig. **6a,b**). At 48 and 72 hpi, unlike in the Cy3 and LDH+Cy3-sprayed pea leaves, Cy3 signals were visible within conidia and primary hyphae present on dsRNA-Cy3-sprayed leaves and conidia present on dsRNA-Cy3-LDH-sprayed leaves (Fig. **6c,d**), indicating uptake of Cy3-dsRNA by the fungus. It is important to note that the fluorescence signal was not consistently observed across all conidia, possibly due to the limited Cy3 labelling efficiency (∼40%). Despite this variability, a consistent inhibitory effect on fungal growth was observed in dsRNA-Cy3 and dsRNA-Cy3-LDH-sprayed leaves when compared to Cy3 alone and Cy3+LDH, with this effect being more pronounced in the dsRNA-Cy3-LDH-sprayed leaves (Fig. **6d**). Further, although a slight delay in fungal growth was observed early on (24 hpi) in the LDH+Cy3-sprayed leaves compared to the Cy3-sprayed controls (Fig. **6b**), this effect was transient, with fungal growth resuming on these leaves at later time points (Fig. **6c,d**).

**Fig. 6.**
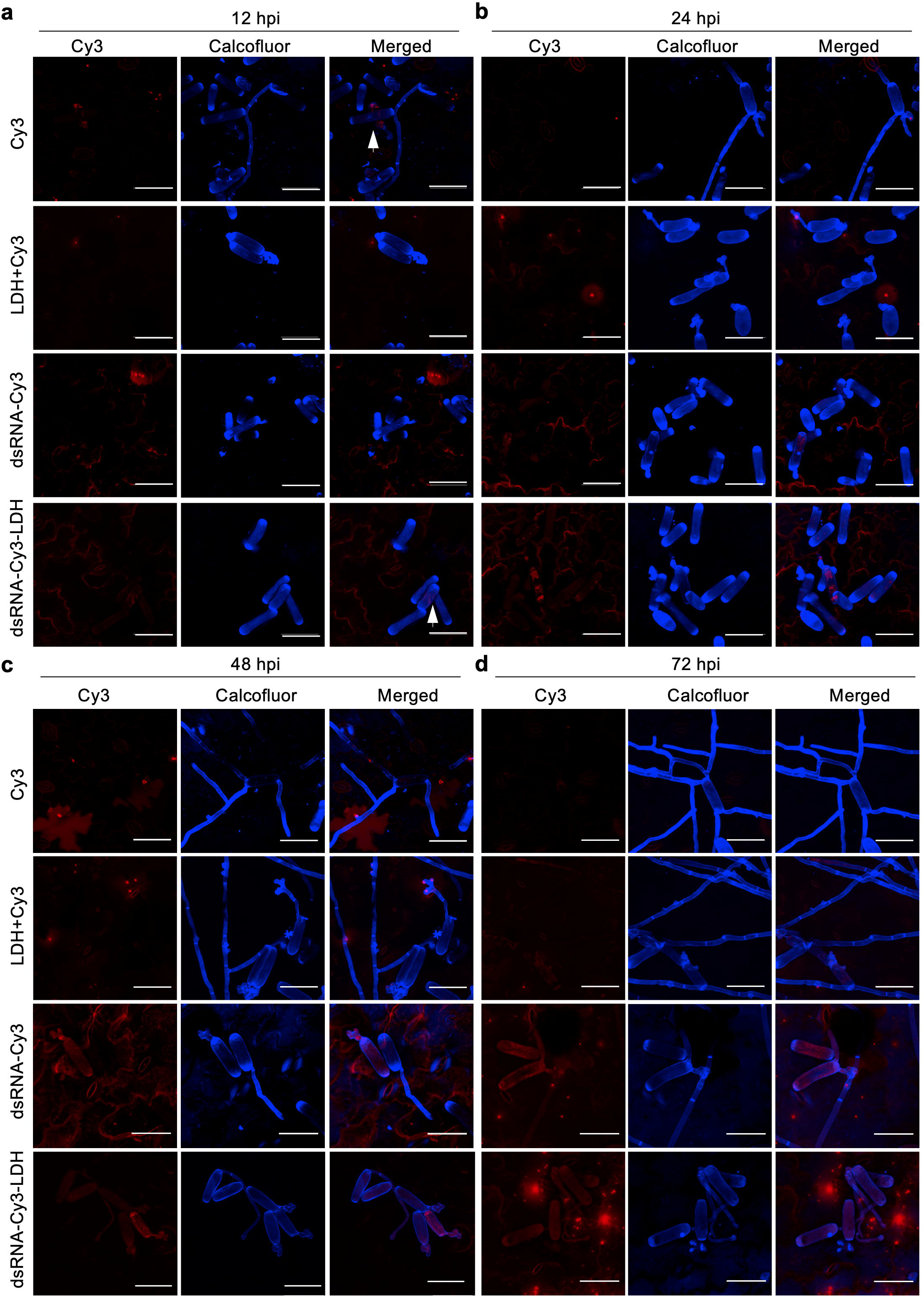
Uptake of dsRNA loaded on MgFe-LDH by PM conidia from pea leaves. (a-d) Representative confocal images showing uptake of dsRNA-Cy3, dsRNA-Cy3-LDH, LDH+Cy3, or Cy3 alone into fungal conidia and/or hyphae at 12-72 hours post-inoculation (hpi) with *E. pisi.* The fungus is stained with calcofluor and shown in blue, and the Cy3 fluorescence signal is shown in red; Scale *=* 50 μm

To determine whether systemic movement of the Ef-dsRNA is responsible for the fungal growth inhibition in unsprayed leaves, pea leaves were sprayed with Ef-dsRNA-Cy3 or Ef-dsRNA-Cy3-LDH and placed in the dark for 24 hours. The next day, *E. pisi* conidia were brush-inoculated on the sprayed leaves of a subset of plants. At 15 days post-spray (dps), leaf tissues of sprayed leaves and newly emerged, unsprayed leaves were harvested from inoculated and non-inoculated plants and analyzed using confocal microscopy. A stronger Cy3 signal was detected in sprayed and newly emerged, unsprayed leaves of Ef-dsRNA-Cy3-LDH-treated plants than in dsRNA-Cy3-treated pea plants (Fig. **S7a**), suggesting that LDH enhances the uptake and systemic mobility of dsRNA within the plant. Similarly, a stronger Cy3 signal was observed at fungal haustorial sites in sprayed and newly emerged unsprayed leaves of Ef-dsRNA-Cy3-LDH-treated than dsRNA-Cy3-treated inoculated pea plants (Fig. **S7b**). Additionally, fewer fungal hyphae and conidiophores (asexual reproductive structures) were visible on sprayed and systemic leaves of Ef-dsRNA-Cy3-LDH-treated plants compared to the Ef-dsRNA-Cy3-treated controls (Fig. **S7b**), supporting a role for LDH in mediating stability and long-range delivery of dsRNA.

Taken together, these findings suggest that LDH-facilitated internalization of Ef-dsRNA into pea leaf cells and fungal hyphae contributes to the extended and systemic protection afforded by the Ef-dsRNA-LDH formulation against PM.

### MgFe-LDH nanomaterials are biocompatible and gradually degrade on the leaf surface

To assess biocompatibility of the LDH, seed germination assays were conducted using two LDH concentrations (4 and 10 mg mL⁻¹) in pea, *Arabidopsis, Medicago*, and rice under short-and long-term exposure regimes. No negative effects were observed; rather, *Arabidopsis* and *Medicago* showed enhanced germination and/or radicle growth (Fig. **S8-S10**). Further, 10 mg mL⁻¹ LDH was tested for ROS generation, MDA accumulation, and expression of defence/stress genes in pea (Fig. **S11-S13**). A slight but inconsistent increase in H₂O₂/MDA and expression of stress markers was observed at 1 dps across replicates (Fig. **S11–S13**). These findings suggest that even at high concentrations, MgFe-LDH is non-toxic and does not induce a consistent stress response in pea.

To evaluate whether LDH adhering to the leaf surface naturally degrades over time, we performed SEM imaging of pea leaves sprayed with water (control) or MgFe-LDH at 2 hours and 7 days post-spray (hps; dps). Compared to 2 hps, comparatively fewer particles were visible on the surface of LDH-treated leaves at 7 dps, especially near stomata (Fig. **7a****; S14**), implying that the LDH degrades under ambient conditions over time. SEM-EDS analysis revealed a significant decrease in Mg and Fe atom count from 2 hps to 7 dps (Fig. **7b**), confirming LDH breakdown. Chlorophyll content was significantly higher at 7 dps (Fig. **7c**), whereas no significant difference in plant height was observed on LDH treatment (Fig. **7d**).

**Fig. 7.**
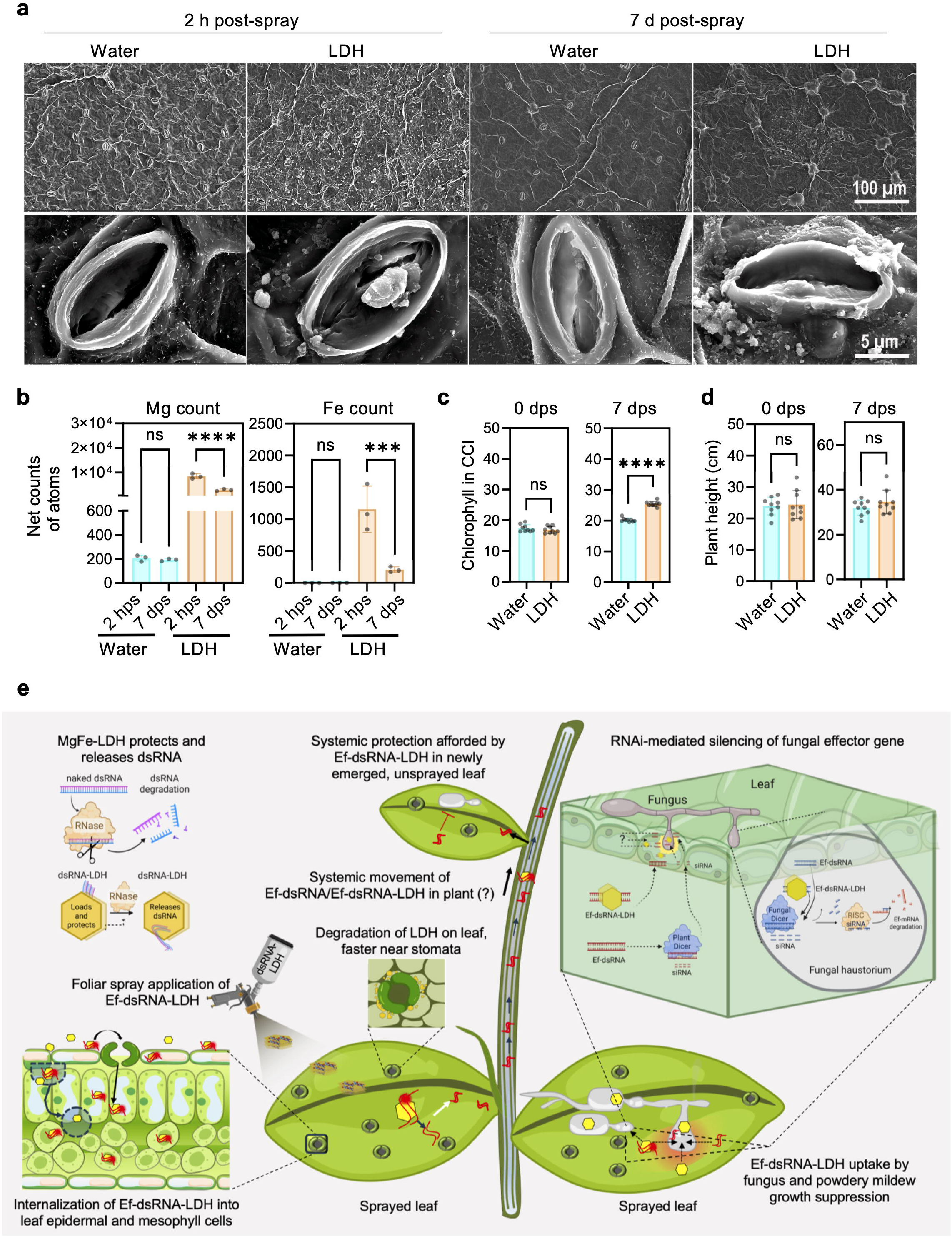
MgFe-LDH degradation on plant leaf surface and effect on plant chlorophyll content and height. (a) Representative SEM images of pea leaf samples captured at 2 hours and 7 days post spray (hps/dps) application of LDH suspension (4mg/mL) or water. The images show the gradual degradation of LDH on the leaf surface (upper panel) and around the stomata (lower panel) at 7 dps. (b) Quantification of Mg and Fe net count of atoms through EDX. Data represent the mean (± SD) of atom counts from three images and three independent experiments. (c) Chlorophyll content index (CCI) measurement. Data represent the mean (± SD) of chlorophyll content from at least 10 regions of nine leaves each and three independent experiments. (d) Plant height at day 0 and day 7 post-LDH or water spray treatment. Data represent the mean (± SD) of 9 plants from three independent experiments. Statistical significance in (b) was computed using Ordinary one-way ANOVA along with Sidak’s multiple comparisons test and in (c-d) via an unpaired t-test (α < 0.05); *** = P ≤ 0.001; **** = P ≤ 0.0001; ns= not significant (e) Schematic showing how fungal effector dsRNA delivery via MgFe-LDH affords prolonged local and systemic RNAi-based protection against powdery mildew in pea. MgFe-LDH protects dsRNA from degradation and facilitates its gradual release through anion exchange or environmental degradation that occurs more rapidly around stomata. Upon foliar spray, Ef-dsRNA or the Ef-dsRNA-LDH complex enters the leaf tissue via stomatal openings or an unknown mechanism and is subsequently taken up as intact dsRNA-LDH complexes, dsRNA alone, and/or processed siRNAs by the fungus through haustorial interfaces, ultimately leading to fungal gene silencing. The released dsRNA (or processed siRNAs) also translocate to newly emerged, unsprayed leaves, offering systemic protection against the disease.

## Discussion

This study presents the first use of dsRNA-LDH technology against a biotrophic fungal pathogen, expanding its proven application against viruses, insects, and necrotrophic fungi (Mitter *et al*., 2017; Jain *et al*., 2022; Niño-Sánchez *et al*., 2022). As summarized in Fig. **7e**, we demonstrate that MgFe-LDH nanocarriers effectively deliver *Erysiphe pisi* effector-targeting dsRNA into plant and fungal cells when topically applied, enhancing and prolonging RNAi-based silencing of the target gene and, consequently, PM growth on pea, an economically important legume crop. Importantly, we show that MgFe-LDH enters fungal structures from infected plant leaf tissues, facilitating the delivery of intact dsRNA into target tissues for efficient gene silencing.

Functionally, the Ef-dsRNA-LDH complex significantly outperformed the naked Ef-dsRNA in reducing *Ef* transcript levels and PM disease symptoms on pea leaves up to 15 dpi. A single spray afforded fungal protection for at least 15 days in sprayed and newly emerged, unsprayed pea leaves when challenged with *E. pisi* (Fig. **1**). These observations indicate that the Ef-dsRNA-LDH complex or the released Ef-dsRNA was effectively internalized within plant and fungal cells of sprayed leaves and systemically translocated within the plant, enhancing RNAi efficiency and enabling disease protection beyond the initial application site. Internalization of 25-60 nm MgAl-LDH nanoparticles (or their modified forms) was previously reported in cells of various plant tissues, including tomato pollen (Yong *et al*., 2021), *N. benthamiana* leaves (Yong *et al*., 2022), *Arabidopsis* roots (Bao *et al*., 2016), and flower pedicels (Molesini *et al*., 2022); however, direct foliar uptake of MgFe-LDH has not been explored. Further, while internalization of diverse nanomaterials into fungal hyphae has been demonstrated, largely in *in vitro* systems (Muse *et al*., 2014; Chen *et al*., 2022; Niño-Sánchez *et al*., 2022; Mukherjee *et al*., 2024), uptake of LDH by fungi from plant tissues has not been reported. This dual internalization is pivotal for efficient and targeted delivery of dsRNA in RNAi-based plant protection strategies. Here, we demonstrate the internalization of 60-150 nm FITC-MgFe-LDH into leaf epidermal and mesophyll cells of multiple plant species, including pea, *N. benthamiana*, and rice, within an hour of spray application (Fig. **3**; Fig. **S5,6**). Notably, the nanocarrier colocalizes with chloroplasts of mesophyll cells as previously reported for MgAl-LDH (Yong *et al*., 2022). We also show that FITC-LDH enters *E. pisi* hyphae from infected pea leaves, likely through fungal haustoria, specialized biotrophic interfaces through which PMs acquire host-derived nutrients (Mapuranga *et al*., 2022). 3D confocal reconstructions confirmed the presence of FITC-LDH within the hyphal network, suggestive of LDH movement through internal fungal channels (Fig. **4**). The strong adherence of FITC-LDH to the leaf surface, even after washing, may promote its internalization into plant cells and fungal hyphae (Fig. **3a**). Yong *et al*. (2025) recently showed that lysozyme-functionalized MgAl-LDH (30–40 nm) enters *N. benthamiana* leaves via an endocytosis-mediated cellular uptake. While the precise mechanism of MgFe-LDH uptake remains unclear, our findings underscore the capacity of these nanomaterials to cross both plant and fungal cellular barriers.

We further demonstrate that co-delivery with MgFe-LDH improves the Ef-dsRNA stability and uptake. Firstly, Ef-dsRNA was detected on Ef-dsRNA-LDH-sprayed leaves at 15 days post PM challenge as opposed to only 7 days on Ef-dsRNA-sprayed leaves (Fig. **2d**). Secondly, dsRNA-Cy3 signal was detected in leaf cells and fungal hyphae of sprayed and systemic leaves up to 15 days after Ef-dsRNA-LDH application or 14 days post *E. pisi* inoculation (Fig. **S7**). MgAl-LDH similarly enhanced the stability of viral gene-specific dsRNA, providing prolonged local and systemic protection against cucumber mosaic virus in *N. tabacum* plants (Mitter *et al*., 2017). Surprisingly, at the early infection stages (12–72 hpi), no noticeable difference in dsRNA-Cy3 signal was observed in fungal cells when the Ef-dsRNA-Cy3 was delivered with LDH, possibly due to the greater immediate availability of free dsRNA in pea leaves treated with dsRNA-Cy3 alone (Fig. **6**). Consequently, fungal growth was similarly inhibited in both treatments at early infection time points.

Developing biocompatible nanocarriers is essential for advancing sustainable crop protection. Jain *et al*. (2022) showed that 1.5 µg µL⁻¹ MgFe-LDH treatment did not activate abiotic stress-related markers or physiological stress in plants. Likewise, Wang *et al*. (2024) showed that exposure to 40 µg mL⁻¹ MgFe-LDH did not increase apoptosis in neural progenitor cells compared to controls. In our study, higher MgFe-LDH concentrations were required to achieve partial (1:20) and complete (1:50) dsRNA loading, likely due to the lower surface charge and consequently weaker electrostatic potential of the nanomaterial, a property also reported by Wang et al. (2024). Therefore, we assessed the phytotoxicity of LDH at a higher concentration. We found that application of 10 mg mL^-1^ MgFe-LDH did not impair seed germination or early vegetative growth across several plant species (Fig. **S8-10**). Instead, a positive effect on seed germination was observed in Arabidopsis and *M. truncatula* (Fig. **S9-10**), consistent with a previous study in cucumber (Wu *et al*., 2024). In addition, the MgFe-LDH-treated plants exhibited a higher chlorophyll content (Fig. **7c**), an effect potentially attributed to the gradual release of essential micronutrients (Mg and Fe) from the LDH matrix, as it naturally degrades on the leaf surface over time (Fig. **7a,b**). Mg contributes to chlorophyll biosynthesis (Ogunyemi *et al*., 2023), whereas Fe plays a vital role in photosynthetic electron transport and enzyme activity (Ullah *et al*., 2024). Notably, we also observed that the MgFe-LDH present on the leaf surface degrades faster around stomatal regions (Fig. **S14**), likely due to rapid gaseous exchange and pH fluctuations in the stomatal microenvironment (Roelfsema & Hedrich, 2002). Further, the dsRNA-MgFe-LDH complex exhibits enhanced structural stability at a higher temperature (25 °C) than previously reported (Wu *et al*., 2024), with the formulation remaining stable for over 30 days under these conditions (Fig. **1d**), a valuable feature for field applications.

Collectively, our findings highlight MgFe-LDH as an environmentally safe and effective platform for delivering dsRNA to control biotrophic fungal pathogens. Future research should explore multi-target formulations and field scalability. Further, understanding whether LDH can translocate through the vascular system will help resolve whether intact dsRNA or processed siRNAs reach systemic tissues.

## Materials and Methods

### LDH synthesis and characterization

MgFe-LDH nanomaterials were synthesized via a co-precipitation method, modified from Jain *et al*. (2022) and Xu *et al*. (2006). Briefly, a 0.4 M NaOH (Sigma-Aldrich, MO, USA) solution was prepared by dissolving 2.88 g of NaOH pellets in 180 mL of diethylpyrocarbonate (DEPC)-treated water by continuous stirring at 600 rpm in an ice bath for 5 min. To this solution, 5.49 g of MgCl_2_·6H_2_O (Sigma-Aldrich) was added, followed by 2.43 g of FeCl_3_·6H_2_O (Sigma-Aldrich) at a ratio of 3:1 with continuous stirring until the solution appeared dark orange, indicating the onset of precipitation. The pH of the solution was maintained between 9.0-10 with 2 mM NaOH. The suspension was stirred continuously at 600 rpm for 12 hours at 4°C. The resulting precipitate was centrifuged at 3078 rcf for 15 min at RT. The pellet was washed twice with DEPC-treated water, resuspended in 20 mL DEPC-treated water, transferred to a glass Petri dish, dried at 60°C for 12–15 hours, and ground into a fine powder. The synthesized nanomaterial was characterized using various techniques (Methods **S1**).

### Plant and fungal material

Pea (*Pisum sativum*) cv. AP3 plants were cultivated in a controlled environment growth chamber (Conviron, Manitoba, Canada) under the following conditions: 22 °C, a 16-hour light/8-hour dark photoperiod, 70% relative humidity, and photosynthetically active radiation (PAR) of 170 μmol m⁻² s⁻¹. Pure cultures of *Erysiphe pisi* (Palampur-1 isolate) were maintained on AP3 pea plants grown in a separate chamber under identical conditions.

### dsRNA synthesis and LDH loading

The dsRNA targeting *E. pisi CSEP001* (*Ef*) and *GFP* were synthesized using the MEGAscript® in vitro transcription kit (Invitrogen, California, USA) as previously described (Sharma *et al*., 2019). For LDH loading, 1 µg of Ef-dsRNA was mixed with varying concentrations of LDH in ratios ranging from 1:1 to 1:50. The complexes were incubated at 28 °C at 180 rpm for 30 min with gentle agitation. The efficiency of dsRNA loading was assessed via the degree of retention of the dsRNA-LDH complex in the well of a 2% agarose gel. The fluorescence intensities in the sample well and the corresponding gel lanes were quantified using ImageJ (version 1.54f) (https://youtu.be/wg5xAbS6iTQ).

Size and surface charge of the LDH were analyzed before and after dsRNA loading (fully loaded-1:50) in a Dynamic light scattering (DLS) machine (Malvern Nano Zetasizer or nano ZS) at 25°C and 90° angle. The hydrodynamic size (Z-average) and surface charge (zeta potential) for three independently synthesized batches of LDH were recorded before and after dsRNA loading and plotted using OriginPro 2024b (64-bit), v10.1.5.132.

### Stability of LDH-bound dsRNA and temperature-dependent dsRNA release

To test the integrity of the LDH-bound dsRNA, 20 μL reactions containing 1 µg of dsRNA or fully loaded dsRNA-LDH complex (1:50 ratio) and 1 μL of RNase A (2 ng/μL) were incubated for 10 minutes at 37°C. Following incubation, RNase-A was inactivated by adding 0.5 μL of Takara RNase Inhibitor (40 U/µL). The dsRNA was released from the dsRNA-LDH complex by treating the reaction mixture with 10 μL of release buffer at pH 3 (Mitter *et al*., 2017). All mixtures were heated at 65°C for 2 min and loaded on a 2% agarose gel.

To evaluate dsRNA release from LDH at different temperatures, 10 μg of dsRNA was loaded on 500 μg of LDH, and the mixture was brought to a final volume of 100 μL using nuclease-free water (NFW). As a control, the same amount of dsRNA (10 μg) was dissolved separately in 100 μL of NFW. Three aliquots (25 μL each) of each mixture were incubated at 4 °C or 25 °C. At 7, 15, and 30-days post-incubation, 15 μL (corresponding to ∼1–1.5 μg dsRNA) of each sample was analysed on a 2% agarose gel.

### dsRNA-LDH crop protection assays

For the crop protection assays, a 1:20 Ef-dsRNA-LDH mass ratio was used instead of the complete loading ratio of 1:50, to ensure effective Ef-dsRNA delivery at minimal LDH concentration, reduce potential long-term phytotoxicity, and provide some free dsRNA for immediate action after spraying. Leaves of 20-day-old pea plants were sprayed with GFP-dsRNA (200 ng/μL), Ef-dsRNA (200 ng/μL), LDH (4 μg/μL), GFP-dsRNA-LDH (1:20), or Ef-dsRNA-LDH (1:20) prepared in 0.01% Tween 20. Each treatment included six leaves/plant from a total of five plants. One day later, the sprayed leaves were inoculated with *E. pisi* at a dose of ∼10,000 conidia/mL, and PM disease symptoms were quantified at 7, 10, and 15 days post-inoculation (dpi) using ImageJ (Ray & Chandran, 2024).

To test whether Ef-dsRNA-LDH offers prolonged and systemic protection, leaves of 20-day-old pea plants were sprayed with water, Ef-dsRNA (200 ng/μL), LDH (4 μg/μL), or Ef-dsRNA-LDH (1:20) prepared in 0.01% Tween 20. The treated plants were challenged with *E. pisi* (∼10,000 conidia/mL) 15 days post-spray. At 10 dpi, the initially sprayed leaves and the newly emerged unsprayed leaves were harvested, and PM disease symptoms were quantified.

### RNA Isolation, cDNA synthesis, and reverse transcription-quantitative PCR (RT-qPCR)

Total RNA was isolated from ∼100 mg pea leaves using the Nucleospin RNA Plant kit (Macherey-Nagel, Germany) with on-column DNase treatment per the manufacturer’s protocol. First-strand cDNA was synthesized from 1.5 µg total RNA utilizing the Prime-Script cDNA synthesis kit (Takara, Japan) per the manufacturer’s instructions. The resulting cDNA was diluted two-fold and employed for RT-qPCR using the 5X TB Green Premix qPCR mix (Takara-Bio Inc.) in a QuantStudio™ 6 Flex Real-Time PCR system (Applied Biosystems, USA). Four to six independent biological replicates were analysed from each experiment. Relative expression values of target genes (*EpEf* and *Eptub2*) normalized to the geometric mean (GM) of two plant reference genes [*Pisum sativum tubulin beta-3* (*Pstub3*; NM_001427574.1) and *Ps Protein Phosphatase 2A* (*PsPP2A*; NM_001427333.1)] were calculated using LinRegPCR [v.2021.2; (Ruijter *et al*., 2009)]. The specificity of the amplified products was confirmed through melt curve analysis. Primer sequences are provided in **Table S1**.

### RNA Dot-Blot

Total RNA from sprayed leaf samples was pooled and concentrated to ∼2 μg/μL. Synthesized dsRNA (∼4 ng/μL) was used as a positive control, and total RNA (∼2 μg/μL) from unsprayed leaves was used as a negative control. 5 μL of each RNA was spotted on a nylon membrane (Roche, Cat No. 11209299001), activated with water, followed by 20x SSC. The membrane was dried for 10 min, followed by UV cross-linking at 22J for 2 min in a UV-Cross linker (Cleaver Scientific, CL-508). The cross-linked nylon membrane was pre-hybridized in Prehyb buffer for 1-2 hours at 50°C. The DIG-labelled PCR-probes were synthesized from cDNA derived from the *Ef*-dsRNA per the manufacturer’s protocol (Roche, Cat. No. 11636090910). The prehyb solution was replaced with hybridization solution containing the Dig-labelled PCR-probe and incubated at 50°C at 10 rpm overnight. The membrane was processed per the manufacturer’s protocol for the DIG detection kit (Roche, Cat. No. 11363514910) and analyzed in a Bio-Rad ChemiDoc MP Imaging System.

### LDH adherence and internalization into pea leaves

MgFe-LDH was fluorescently labelled with Fluorescein isothiocyanate (FITC; Cat. No. 46425, Thermo Fisher Scientific), following the protocol described by Yong *et al*. (2022). Briefly, 25 μL of FITC (10 mg/mL in dimethylformamide, DMF) was added to 500 μL of MgFe-LDH (10 mg/mL in DEPC-treated water) in a 2 mL microcentrifuge tube. Subsequently, 50 μL of NaOH (pH 11) was added. The tube was wrapped in aluminium foil and incubated at 26 °C in a shaker incubator for 1 hour. After incubation, the mixture was centrifuged at 15,000 rpm for 1 min at room temperature (RT), and the supernatant was discarded. The resulting pellet was washed with 500 μL of 70% ethanol and centrifuged under the same conditions. The pellet was washed twice with 1 mL DEPC-treated water, with each step followed by centrifugation. Finally, the supernatant was removed, and the FITC-LDH complex (1:20 ratio) was resuspended in 500 μL NFW.

To test adherence, FITC (10 μg/mL) and FITC-labelled LDH (FITC-LDH 1:20) were sprayed on leaves of 20-day-old pea plants. Following 1 hour dark incubation, both sets of leaves were washed with autoclaved Milli-Q water and blot-dried with Kim-wipes. Leaves were visualized under a handheld UV lamp (UVP Blak-Ray B-100AP, 365 nm, long-wave UV). Images were captured using a smartphone camera before wash, and at 1 and 12 hours post-wash.

For LDH internalization studies, the FITC-LDH-sprayed leaves (1 hour post-spray) were washed thoroughly and blot-dried with Kim-wipes. The leaves were sectioned into 0.5 cm^2^ square pieces and visualized under the 40× oil objective of a Leica SP8 CLSM with the following laser settings: 5%-Argon laser set at 50% gain with HyD detector for FITC-green channel (λex/λem): 490 nm/500–520 nm; chloroplast autofluorescence-purple channel (λex/λem): 490 nm/610–750 nm. Z-stack images at a step size of 0.5 μm were captured, and optical slices representing the epidermis (leaf upper surface) and mesophyll layers (cell layers characterized by high autofluorescence) were merged separately. The images were captured at two magnifications, 0.75x and 5x, and processed using the Leica Application Suite X (LAS-X, v.-3.5.7.23225). Colocalization of FITC fluorescence with chloroplast autofluorescence was analyzed using line scans. The fluorescence intensity along each line was measured, and the raw intensity data were plotted using OriginPro 2024b, as previously described (Kreft *et al*., 2010; Yong *et al*., 2022).

### LDH uptake into PM fungus from pea leaves

FITC (10 μg/mL) and FITC-labelled LDH (1:20) were sprayed on leaves of 20-day-old pea plants. Following 1 hour dark incubation, both sets of leaves were washed with autoclaved Milli-Q water and blot-dried with Kim-wipes. The sprayed leaves were inoculated by brushing *E. pisi* conidia from 12 dpi pea leaves and harvested at different hours post-inoculation (hpi) for confocal microscopy. To visualize the fungal structures, infected leaf sections (∼0.5 cm^2^) were stained with one drop of Calcofluor White (Sigma Aldrich, 18909-100ML-F) and one drop of 10% potassium hydroxide and incubated for 5 min at RT. The leaf sections were imaged under the 40× or 63× oil objective of a Leica SP8 CLSM with the following laser settings: DAPI laser (405 nm) at 1–2% intensity with the HyD detector (gain: 20–30%) for calcofluor white-blue channel (λ_ex_ = 403 nm, λ_em_ = 433 nm) and 5%-Argon laser, set at 50% gain with HyD detector for FITC-green channel (λ_ex_/λ_em_): 490 nm/500–520 nm. Captured Z-stack images (step-size of 0.35 μm) were processed by the Leica Application Suite X, followed by 3D surface recreation using Imaris x64 v.8.3.0. The reconstructed 3D surface of the fungal body was pseudo-coloured metallic blue with solid fill or 60% transparency.

### dsRNA uptake into pea leaves

The Cy3 labelling kit (Invitrogen, AM1632) was used to fluorescently tag Cy3 to ∼20 µg of dsRNA, following the manufacturer’s instructions. Half of the labelled dsRNA (∼10 µg) was loaded on MgFe-LDH at a 1:50 dsRNA:LDH ratio. Cy3 (1000X dilution), Cy3-labeled dsRNA (dsRNA-Cy3), or dsRNA-Cy3-LDH (1:50) was sprayed on leaves of 20-day-old pea plants. Post-application, the plants were placed in the dark for 1 hour, washed thoroughly with autoclaved Milli-Q water, blot-dried using Kim-wipes, and returned to darkness for 11 hours.

Confocal imaging was performed using a Leica SP8 CLSM under a 40× oil-objective lens with the following laser settings: 5%-Argon laser set at 80% gain with HyD detector for Cy3-red channel (λ_ex_/λ_em_): 550 nm/590, and chloroplast autofluorescence-green channel (λ_ex_/λ_em_): 490 nm/610–750 nm. Z-stack images were acquired at 0.5 µm intervals. The optical slices representing epidermal and mesophyll layers were merged separately as described above. Fluorescence intensity for the Cy3 signal was quantified using ImageJ by applying thresholding to isolate the Cy3 signal and subtract background fluorescence. The signal intensity was calculated as the total pixel count, summed across at least three images from two independent experiments.

### dsRNA uptake into fungal hyphae

Leaves of 20-day-old pea plants were sprayed with dsRNA-Cy3 (200 ng/μL), dsRNA-Cy3-LDH (1:20), LDH+Cy3 (1:20), or a 1000X dilution of Cy3 and placed in the dark for 1 hour. Leaves were washed with autoclaved Milli-Q water, blot-dried, and brush-inoculated with *E. pisi* conidia after 30 min. Leaves were harvested at different time points after inoculation (12, 24, 48, and 72 hpi), sectioned, and stained with Calcofluor White, as previously described. The leaf sections were imaged under the 40× oil-objective lens of a Leica SP8 CLSM with the following laser settings: DAPI laser (405 nm) at 1–2% intensity with the HyD detector (gain: 20–30%) for calcofluor white-blue channel (λex = 403 nm, λem = 433 nm); 5%-Argon laser set at 80% gain with HyD detector for Cy3-red channel (λex/λem): 550 nm/590. The captured Z-stacks (0.35 μm intervals) were processed in the Leica Application Suite X.

### LDH degradation on pea leaves and effect on plant physiology

Leaves of 20-day-old pea plants were sprayed with water or LDH (4 mg/mL or 10 mg/mL) using an airbrush (Zorbes Spirit Air 0.4 mm Multi-Purpose Airbrush). The sprayed leaves were harvested at 2 hours post-spray (hps) and 7 days post-spray (dps), fixed in 40% formaldehyde for 5–10 hours, and serially dehydrated with an ethanol series (10%, 20%, 30%, 50%, and 70%, 10 min each). Leaf sections (∼ 0.5 cm^2^) were immersed in 100% acetone twice for 30 min each (Pathan *et al*., 2008), mounted on aluminium stubs with carbon-coated tape, sputter-coated with gold for 40–50 sec, and imaged using Apreo Volume Scope FEI SEM. LDH degradation was quantified by measuring the decrease in atomic percentages of Mg and Fe between 2 hps and 7 dps using SEM-EDS.

Leaf chlorophyll content was measured on a circular 0.8 cm^2^ leaf area using an Apogee SPAD meter (ME-100). Ten readings were taken per leaf on day 0 (before spray) and 7 days after LDH spray. Plant height was also measured at the same time points.

### Statistics

Statistical analyses were performed using GraphPad Prism software (v.10.3). Supplementary Methods are provided in Methods **S1**.

## Author Contributions

PR, BP and DC planned and designed the research. PR, MB and SS performed experiments and analyzed data. PR, MB and DC wrote the manuscript.

## Acknowledgments

We thank Suraj Tiwari, Madhav Rao and Reena Rani from the Advanced Technology Platform Centre for helping with the SEM and TEM microscopy facility. We thank Sprint Testing Solutions for HRTEM, SAED, XPS, and BET analysis. We are grateful to Dr. Raghavendra Aminedi for assisting with the initial experimental design, Dr. Shaily Tyagi and Dr. Kesiraju Karthik for assisting with the Dot Blot assay. We also thank Suraj Tiwari, Dr. Akriti Sharma, and Dr. Chandan Kumar for help with confocal imaging. We thank Kalaiyazhagi Rajagopal for helping with LDH synthesis and testing its effect on seed germination.

## Data availability

The data that support the findings of this study are available in Figs 1-8 and Supporting Information.

## Supporting Information

**Table S1.** List of RT-qPCR primers

**Figure S1.** MgFe-LDH characterization via TEM, HR-TEM, FE-SEM, XPS, FTIR, XRD, and BET

**Figure S2** TEM images of MgFe-LDH and dsRNA:LDH

**Figure S3.** Effector dsRNA dose selection for powdery mildew inhibition assays

**Figure S4.** Characterization of MgFe-LDH labelled with FITC

**Figure S5.** Uptake of MgFe-LDH in *Nicotiana benthamiana* leaves after spray application

**Figure S6.** Uptake of MgFe-LDH in rice leaves after spray application

**Figure S7.** LDH facilitates systemic movement of dsRNA into newly emerged, unsprayed pea leaves

**Figure S8.** Effect of short-term and long-term exposure of MgFe-LDH on pea seed germination

**Figure S9.** Effect of short-term exposure of MgFe-LDH on germination of Arabidopsis, *Medicago truncatula,* and rice seeds

**Figure S10.** Effect of long-term exposure of MgFe-LDH on germination of Arabidopsis, *Medicago truncatula,* and rice seeds

**Figure S11.** Effect of MgFe-LDH spray application on ROS generation in mature pea plants (Experiment 1)

**Figure S12.** Effect of MgFe-LDH spray application on ROS generation in mature pea plants (Experiment 2)

**Figure S13.** Effect of MgFe-LDH spray application on ROS generation in mature pea plants (Experiment 3)

**Figure S14.** SEM images showing degradation of MgFe-LDH deposited on/adjacent to pea leaf stomata

**Methods S1.** Supplementary Methods

## References

Bao, W., Wang, J., Wang, Q., O’hare, D. & Wan, Y. 2016. Layered double hydroxide nanotransporter for molecule delivery to intact plant cells. Sci Rep, 6, 26738.

Bekele-Alemu, A., Dessalegn-Hora, O., Safawo-Jarso, T. & Ligaba-Osena, A. 2025. Rethinking progress: harmonizing the discourse on genetically modified crops. Front Plant Sci, 16, 1547928.

Bojorquez-Quintal, E., Escalante-Magana, C., Echevarria-Machado, I. & Martinez-Estevez, M. 2017. Aluminum, a friend or foe of higher plants in acid soils. Front Plant Sci, 8, 1767.

Brown, J. K. 2015. Durable resistance of crops to disease: a Darwinian perspective. Annu Rev Phytopathol, 53, 513–39.

Chandran, D., Sharopova, N., Vandenbosch, K. A., Garvin, D. F. & Samac, D. A. 2008. Physiological and molecular characterization of aluminum resistance in *Medicago truncatula*. BMC Plant Biol, 8, 89.

Chen, X., Shi, T., Tang, T., Chen, C., Liang, Y. & Zuo, S. 2022. Nanosheet-facilitated spray delivery of dsRNAs represents a potential tool to control *Rhizoctonia solani* infection. Int J Mol Sci, 23.

Devi, J., Mishra, G. P., Sagar, V., Kaswan, V., Dubey, R. K., Singh, P. M., Sharma, S. K. & Behera, T. K. 2022. Gene-based resistance to Erysiphe species causing powdery mildew disease in peas (*Pisum sativum* L.). Genes (Basel), 13.

Fisher, M. C., Henk, D. A., Briggs, C. J., Brownstein, J. S., Madoff, L. C., Mccraw, S. L. & Gurr, S. J. 2012. Emerging fungal threats to animal, plant, and ecosystem health. Nature, 484, 186–94.

Fondevilla, S. & Rubiales, D. 2012. Powdery mildew control in pea. A review. Agronomy for sustainable development, 32, 401–409.

Gebremichael, D. E., Haile, Z. M., Negrini, F., Sabbadini, S., Capriotti, L., Mezzetti, B. & Baraldi, E. 2021. RNA interference strategies for future management of plant pathogenic fungi: prospects and challenges. Plants (Basel), 10.

Godfray, H. C., Mason-D’croz, D. & Robinson, S. 2016. Food system consequences of a fungal disease epidemic in a major crop. Philos Trans R Soc Lond B Biol Sci, 371.

Hidayati, N., Apriliani, D. R., Helda, Taher, T., Mohadi, R., Elfita & A, L. 2019. Adsorption of congo red using Mg/Fe and Ni/Fe layered double hydroxides. J Physics: Conference Series, 1282, 012075.

Jain, R. G., Fletcher, S. J., Manzie, N., Robinson, K. E., Li, P., Lu, E., Brosnan, C. A., Xu, Z. P. & Mitter, N. 2022. Foliar application of clay-delivered RNA interference for whitefly control. Nat Plants, 8, 535–548.

Jha, A. C., Jamwal, S., Kumar, A. & Singh, P. 2019. Loss assessment caused by economically important pea (*Pisum sativum* L.) Diseases and their management in hills of Doda (Jammu & Kashmir) under field condition. Int J Curr Microbiol App Sci, 8, 170–176.

Kreft, M., Prebil, M., Chowdhury, H. H., Grilc, S., Jensen, J. & Zorec, R. 2010. Analysis of confocal images using variable-width line profiles. Protoplasma, 246, 73–80.

Lefranc-Millot, C. & Teichman-Dubois, V. 2019. Protein from vegetable sources: A focus on pea protein. In: Hayes, M. (ed.) Novel Proteins for Food, Pharmaceuticals and Agriculture: Sources, Applications and Advances. First Edition ed.: John Wiley & Sons Ltd.

Mann, C. W. G., Sawyer, A., Gardiner, D. M., Mitter, N., Carroll, B. J. & Eamens, A. L. 2023. RNA-based control of fungal pathogens in plants. Int J Mol Sci, 24.

Mapuranga, J., Zhang, L., Zhang, N. & Yang, W. 2022. The haustorium: The root of biotrophic fungal pathogens. Front Plant Sci, 13, 963705.

Maria, Naz, I., Khan, R., Alam, S. S., Iqbal, O., Akram, S., Rajput, N. A., Younas, M. U., Qasim, M., Ali, I., Elsalahy, H. H., Iqbal, R., Aljowaie, R. M. & Munir, S. 2024. Unleashing the synergistic effect of promising fungicides: a breakthrough solution for combating powdery mildew in pea plants. Front Microbiol, 15, 1448033.

Mitter, N., Worrall, E. A., Robinson, K. E., Li, P., Jain, R. G., Taochy, C., Fletcher, S. J., Carroll, B. J., Lu, G. Q. & Xu, Z. P. 2017. Clay nanosheets for topical delivery of RNAi for sustained protection against plant viruses. Nat Plants, 3, 16207.

Molesini, B., Pennisi, F., Cressoni, C., Vitulo, N., Dusi, V., Speghini, A. & Pandolfini, T. 2022. Nanovector-mediated exogenous delivery of dsRNA induces silencing of target genes in very young tomato flower buds. Nanoscale Adv, 4, 4542–4553.

Mukherjee, S., Beligala, G., Feng, C. & Marzano, S. Y. 2024. Double-stranded RNA targeting white mold *Sclerotinia sclerotiorum* Argonaute 2 for disease control via spray-induced gene silencing. Phytopathol, 114, 1253–1262.

Muse, E. S., Patel, N. R., Astete, C. E., Damann, K. E. & Sabliov, C. M. 2014. Surface association and uptake of poly(lactic-co-glycolic) acid nanoparticles by *Aspergillus flavus*. Therapeutic Delivery, 5, 1179–1190.

Niño-Sánchez, J., Sambasivam, P. T., Sawyer, A., Hamby, R., Chen, A., Czislowski, E., Li, P., Manzie, N., Gardiner, D. M., Ford, R., Xu, Z. P., Mitter, N. & Jin, H. 2022. BioClay prolongs RNA interference-mediated crop protection against *Botrytis cinerea*. J Integr Plant Biol, 64, 2187–2198.

Ogunyemi, S. O., Abdallah, Y., Ibrahim, E., Zhang, Y., Bi, J., Wang, F., Ahmed, T., Alkhalifah, D. H. M., Hozzein, W. N., Yan, C., Li, B. & Xu, L. 2023. Bacteriophage-mediated biosynthesis of MnO(2)NPs and MgONPs and their role in the protection of plants from bacterial pathogens. Front Microbiol, 14, 1193206.

Olivier, F. C. & Annandale, J. G. 1998. Thermal time requirements for the development of green pea (*Pisum sativum* L.). Field Crops Research, 56, 301–307.

Ons, L., Bylemans, D., Thevissen, K. & Cammue, B. P. A. 2020. Combining biocontrol agents with chemical fungicides for integrated plant fungal disease control. Microorganisms, 8.

Ouyang, Y., Xia, Y., Tang, X., Qin, L. & Xia, S. 2025. Trans-Kingdom sRNA silencing in *Sclerotinia sclerotiorum* for crop fungal disease management. Pathogens, 14.

Pathan, A. K., Bond, J. & Gaskin, R. E. 2008. Sample preparation for scanning electron microscopy of plant surfaces—Horses for courses. Micron, 39, 1049–1061.

Quilez-Molina, A. I., Nino Sanchez, J. & Merino, D. 2024. The role of polymers in enabling RNAi-based technology for sustainable pest management. Nat Commun, 15, 9158.

Ray, P. & Chandran, D. 2024. Spray inoculation and image analysis-based quantification of powdery mildew disease severity on pea leaves. MethodsX, 13, 102980.

Ray, P., Sahu, D., Aminedi, R. & Chandran, D. 2022. Concepts and considerations for enhancing RNAi efficiency in phytopathogenic fungi for RNAi-based crop protection using nanocarrier-mediated dsRNA delivery systems. Front Fungal Biol, 3, 977502.

Roelfsema, M. R. G. & Hedrich, R. 2002. Studying guard cells in the intact plant: modulation of stomatal movement by apoplastic factors. New Phytol, 153, 425–431.

Ruijter, J. M., Ramakers, C., Hoogaars, W. M., Karlen, Y., Bakker, O., Van Den Hoff, M. J. & Moorman, A. F. 2009. Amplification efficiency: linking baseline and bias in the analysis of quantitative PCR data. Nucleic Acids Res, 37, e45.

Secic, E. & Kogel, K. H. 2021. Requirements for fungal uptake of dsRNA and gene silencing in RNAi-based crop protection strategies. Curr Opin Biotechnol, 70, 136–142.

Sharma, G., Aminedi, R., Saxena, D., Gupta, A., Banerjee, P., Jain, D. & Chandran, D. 2019. Effector mining from the *Erysiphe pisi* haustorial transcriptome identifies novel candidates involved in pea powdery mildew pathogenesis. Mol Plant Pathol, 20, 1506–1522.

Silva Neto, L. D., Anchieta, C. G., Duarte, J. L. S., Meili, L. & Freire, J. T. 2021. Effect of drying on the fabrication of MgAl layered double hydroxides. ACS Omega, 6, 21819–21829.

Singha Roy, A., Kesavan Pillai, S. & Ray, S. S. 2022. Layered Double Hydroxides for sustainable agriculture and environment: An overview. ACS Omega, 7, 20428–20440.

Stukenbrock, E. & Gurr, S. 2023. Address the growing urgency of fungal disease in crops. Nature, 617, 31–34.

Taning, C. N., Arpaia, S., Christiaens, O., Dietz-Pfeilstetter, A., Jones, H., Mezzetti, B., Sabbadini, S., Sorteberg, H. G., Sweet, J., Ventura, V. & Smagghe, G. 2020. RNA-based biocontrol compounds: current status and perspectives to reach the market. Pest Manag Sci, 76, 841–845.

Ullah, J., Gul, A., Khan, I., Shehzad, J., Kausar, R., Ahmed, M. S., Batool, S., Hasan, M., Ghorbanpour, M. & Mustafa, G. 2024. Green synthesized iron oxide nanoparticles as a potential regulator of callus growth, plant physiology, antioxidative and microbial contamination in *Oryza sativa* L. BMC Plant Biol, 24, 939.

Venu, E., Ramya, A., Babu, P. L., Srinivas, B., Kumar, S., Reddy, N. K., Babu, Y. M., Majumdar, A. & Manik, S. 2024. Exogenous dsRNA-mediated RNAi: Mechanisms, applications, delivery methods and challenges in the induction of viral disease resistance in plants. Viruses, 17.

Vielba-Fernandez, A., Polonio, A., Ruiz-Jimenez, L., De Vicente, A., Perez-Garcia, A. & Fernandez-Ortuno, D. 2020. Fungicide resistance in powdery mildew fungi. Microorganisms, 8.

Wan, N. F., Fu, L., Dainese, M., Kiaer, L. P., Hu, Y. Q., Xin, F., Goulson, D., Woodcock, B. A., Vanbergen, A. J., Spurgeon, D. J., Shen, S. & Scherber, C. 2025. Pesticides have negative effects on non-target organisms. Nat Commun, 16, 1360.

Wang, Z., Bai, Y., Xu, X., Zhu, Y., He, X., Huang, R., Yu, L., Huang, R., Cheng, L. & Zhu, R. 2024. Two-Way Regulation of MgFe-LDH on ESC cell fate via temporal selective activation of different cell membrane receptors. Advanced Functional Materials, 34.

Wu, H., Wan, X., Niu, J., Cao, Y., Wang, S., Zhang, Y., Guo, Y., Xu, H., Xue, X., Yao, J., Zhu, C., Li, Y., Li, Q., Lu, T., Yu, H. & Jiang, W. 2024. Enhancing iron content and growth of cucumber seedlings with MgFe-LDHs under low-temperature stress. J Nanobiotechnology, 22, 268.

Xu, Z. P., Stevenson, G. S., Lu, C.-Q., Lu, G.-Q., Bartlett, P. F. & Gray, P. P. 2006. Stable suspension of Layered Double Hydroxide nanoparticles in aqueous solution. J American Chem Soc, 128, 36–37.

Yong, J., Wu, M., Zhang, R., Bi, S., Mann, C. W. G., Mitter, N., Carroll, B. J. & Xu, Z. P. 2022. Clay nanoparticles efficiently deliver small interfering RNA to intact plant leaf cells. Plant Physiol, 190, 2187–2202.

Yong, J., Xu, W., Wu, M., Zhang, R., Mann, C. W. G., Liu, G., Brosnan, C. A., Mitter, N., Carroll, B. J. & Xu, Z. P. 2025. Lysozyme-coated nanoparticles for active uptake and delivery of synthetic RNA and plasmid-encoded genes in plants. Nat Plants, 11, 131–144.

Yong, J., Zhang, R., Bi, S., Li, P., Sun, L., Mitter, N., Carroll, B. J. & Xu, Z. P. 2021. Sheet-like clay nanoparticles deliver RNA into developing pollen to efficiently silence a target gene. Plant Physiol, 187, 886–899.

Zhang, Y., Xu, H. & Lu, S. 2021. Preparation and application of layered double hydroxide nanosheets. RSC Adv, 11, 24254–24281.

