## Supplementary Information for "Effector dsRNA delivery via MgFe-layered double hydroxide nanocarriers confers prolonged protection against powdery mildew in pea"

**Table S1**. List of RT-qPCR primers

| Primer Name | Forward primer sequence (5′-3′) | Reverse primer sequence (5′-3′) |
|---|---|---|
| *Ep*CSEP001 | GACTTACGGAGAACATGGACAC | GGCAATTATGGCGACTCTCTTA |
| *EpBeta-tubulin 2* (*Eptub2*) | ATCTGCCGTCATGTCAGGTG | CAGTGACAGCCCGGAATGAG |
| *Ep18s rRNA* | TCCAGCTGCCTTTGTGTGG | GATGAGGGTTGTTCTGGCAAGC |
| *PsPP2A* | CGGCTGCTGCGTGATAATG | TGGTAGAATATGCTGAATGGCTAGTT |
| *PsFeSOD* | GGAAGCACCACAGAGCTTAT | ATTGTTGAAAGCCGGAAGAATG |
| *PsPRP4A* | CGCACACCAGCCTCTATTATG | GCACTCTGACCACTCACTAAC |
| *PsChitinase* | TCATCTCCAACACCGACAATAC | CACACCGTCCATCGTTTCTAT |
| *PsPAL* | ACAGTAGCAGCAGCCATAAC | CTCATCCAAGTGACTCCCTTTC |
| *PsHSP71.2* | TCTTCGACCGCACAAACTAC | CACTTCTCAACAGGCTCCATAC |
| *Pstubulin beta-3* | TTGGGCGAAAGGACACTATACTG | CAACATCGAGGACCGAGTCA |

*Ep, E. pisi; Ps, P. sativum*

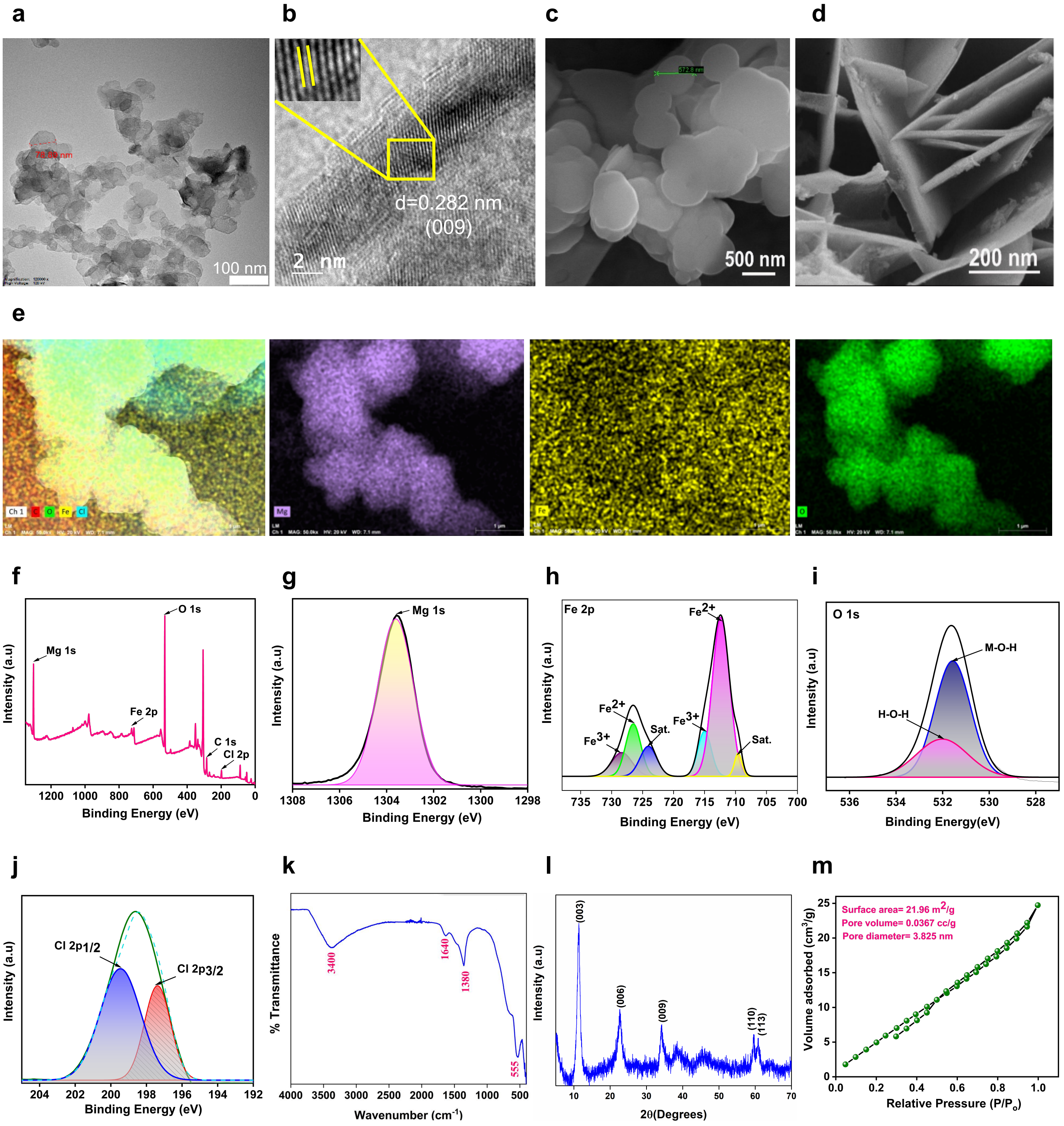


**Fig. S1 Characterization of MgFe-LDH nanomaterial** (a) TEM image (b) HR-TEM image (c) FE-SEM image showing the surface view of the LDH (d) FE-SEM image showing the lateral view of the LDH (e) Elemental mapping of Mg, Fe and O (f) XPS total spectrum of elemental distribution and (g-j) corresponding elemental peaks for Mg 1s, Fe 2p, O 1s and Cl 2p (k) FTIR (l) XRD pattern (m) BET graph using BJH method of adsorption

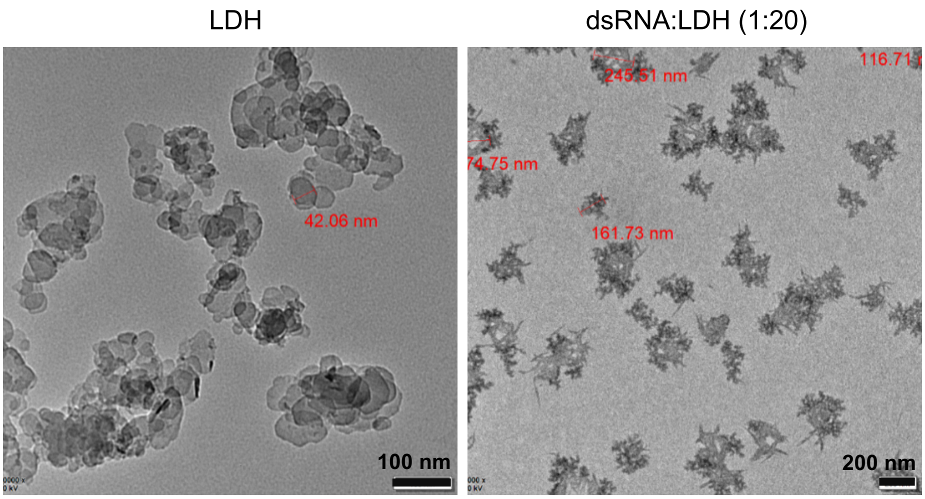


**Fig. S2** TEM images of MgFe-LDH and dsRNA:LDH (1:20)

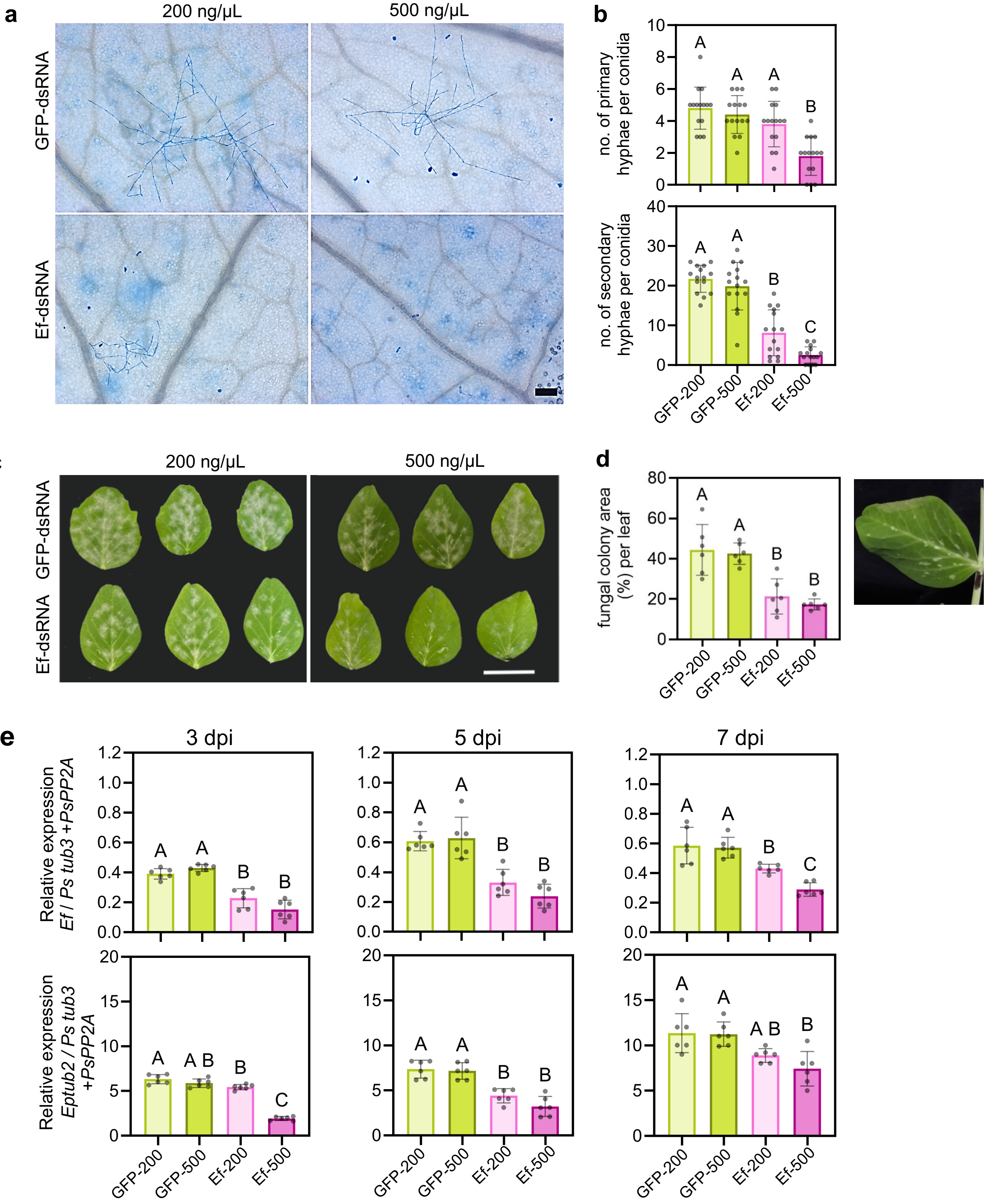


**Fig. S3. Effector dsRNA dose selection for powdery mildew inhibition assays** (a) Representative light microscopy images showing *E. pisi* colony development at 3 days post-inoculation (dpi) on pea leaves sprayed with 200 ng/µL or 500 ng/µL Effector (Ef)-dsRNA or GFP-dsRNA; Scale = 200 µm (b) Bar graphs showing the average number (±SD) of primary hyphae (above) or secondary hyphae (below) per conidium quantified from conidia present in 5 focal areas each from 3 leaves (c) Powdery mildew disease symptoms on pea leaves sprayed with 200 ng/µL or 500 ng/µL Ef-dsRNA or GFP-dsRNA at 7 dpi; Scale = 2 cm (d) Quantification of PM disease symptoms using ImageJ. Data represents mean (±SD) % leaf area covered with disease symptoms at 7 dpi assessed from 6 leaves per treatment. Representative image of a pea leaf showing damage at the leaf margin after 500 ng/µL dsRNA spray (e) Bar graphs showing relative expression of *EpEf* (above) and *Eptub2* (below) normalized to the geometric mean of two endogenous controls, *Pstub3 and PsPP2A,* at 3, 5 and 7 dpi; n=6 leaves per treatment and time point. Statistical significance was computed via One-way ANOVA ($p < 0.0001$) along with Tukey's multiple comparisons test and denoted as uppercase letters.

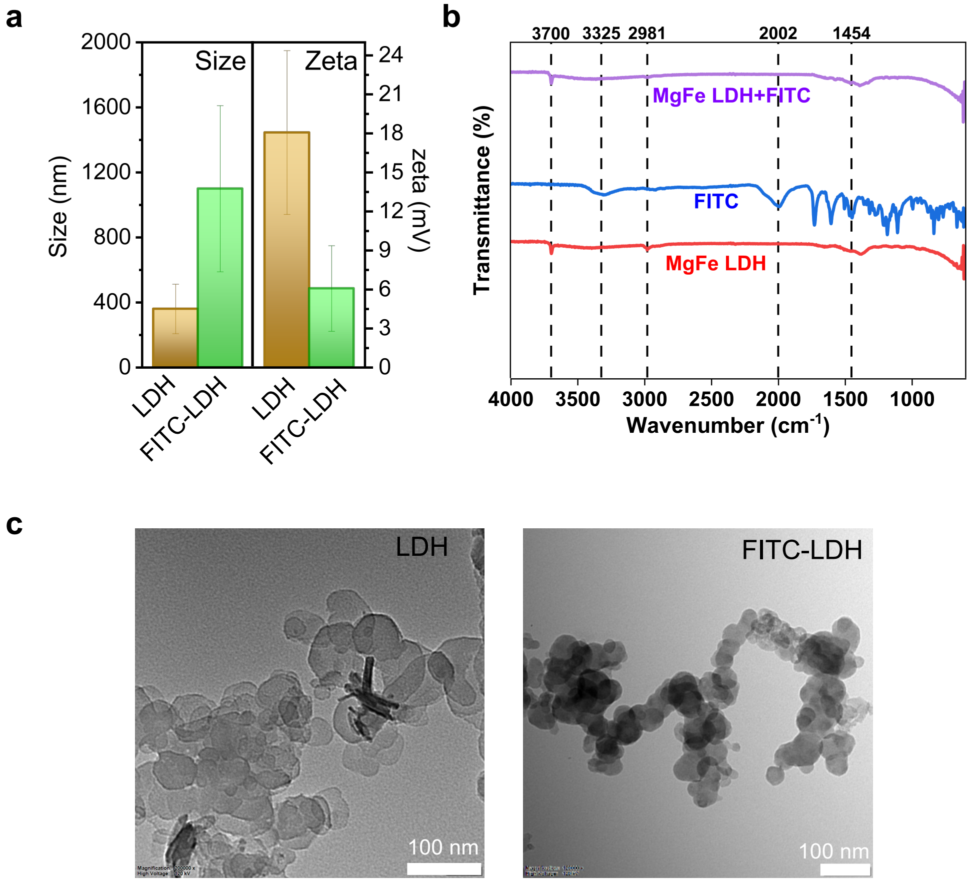


**Fig. S4 Characterization of MgFe-LDH labelled with FITC** (a) Bar graphs showing mean (±SD) of size and surface charge of LDH (0.1 mg/mL) before and after FITC labelling (FITC:LDH = 1:20), analyzed via DLS; n=3 (b) FTIR analysis showing a change in functional groups (at 1454 and 2981 wave number) of LDH post-FITC labeling (c) TEM images of LDH and FITC-LDH

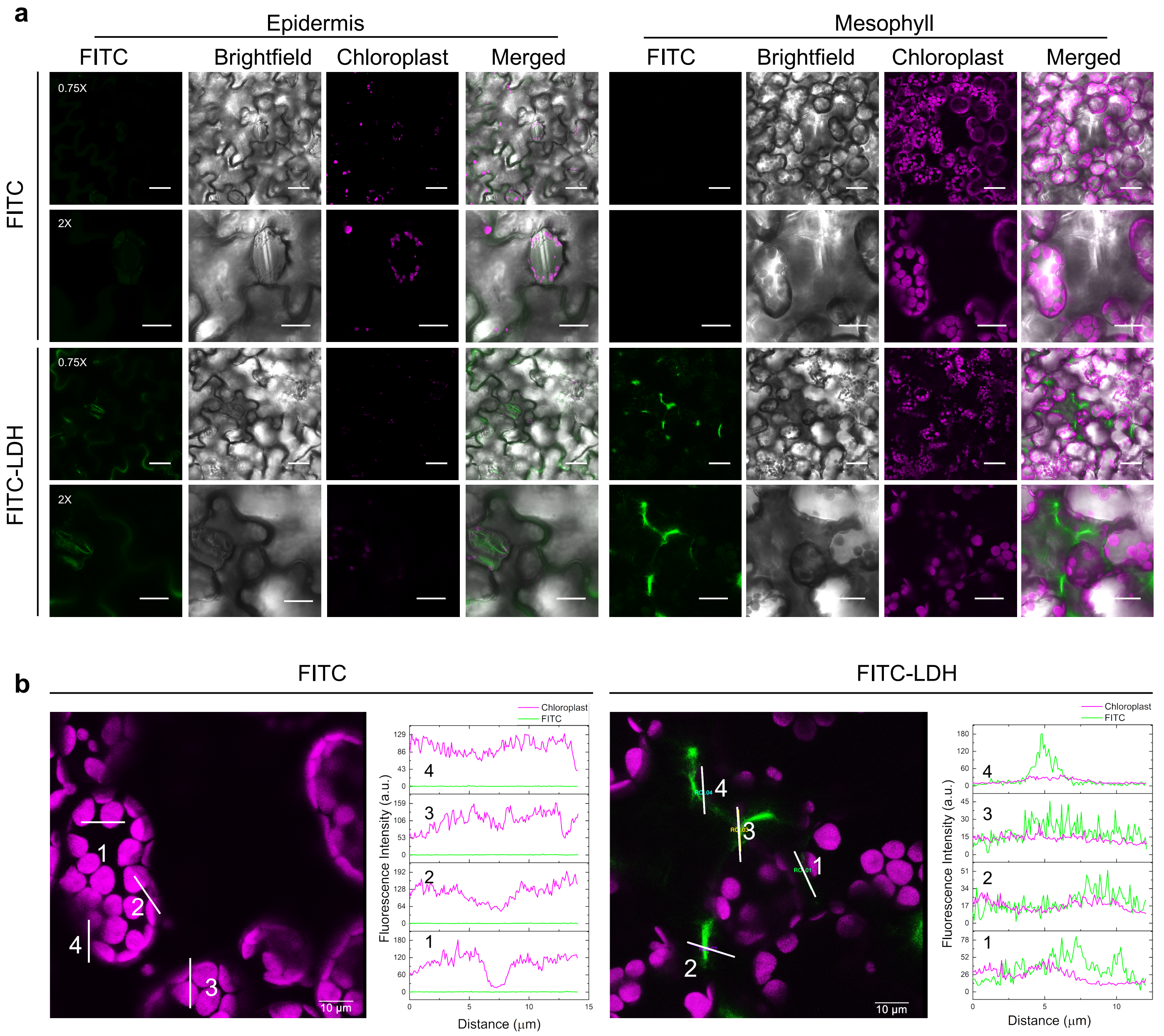


**Fig. S5 Uptake of MgFe-LDH in *Nicotiana benthamiana* leaves after spray application** (a) Representative confocal images showing LDH internalization into epidermal and/or mesophyll cells of *N. benthamiana* leaves. The adaxial surface of *Nicotiana* leaves was sprayed with FITC-LDH (1:20) or FITC alone, rinsed with water after 1 h, and observed immediately under a 63x oil objective of a Leica SP8 confocal microscope at magnification 0.75x (Scale = 40 µm) and magnification 2x (Scale = 15 µm). FITC fluorescence is visible as green, and chloroplast autofluorescence as magenta. (b) Magnified view of the mesophyll cells of FITC and FITC-LDH sprayed *N. benthamiana* leaves along with line scans (white lines 1-4) and corresponding intensities of FITC fluorescence (green) and chloroplast autofluorescence (magenta).

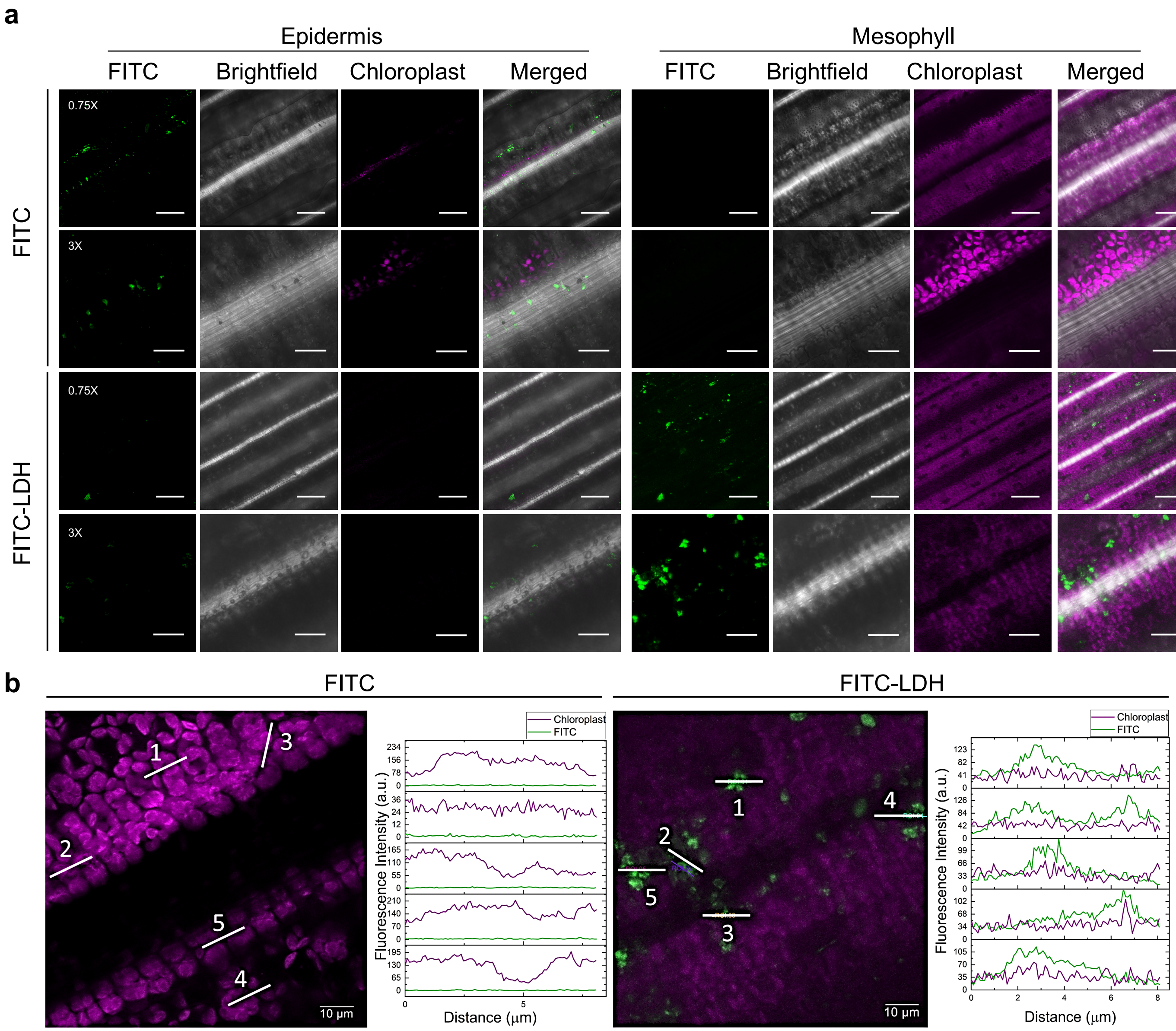


**Fig. S6 Uptake of MgFe-LDH in rice leaves after spray application** (a) Representative confocal images showing LDH internalization into epidermal and/or mesophyll cells of rice leaves. The adaxial surface of rice leaves was sprayed with FITC-LDH (1:20) or FITC alone, rinsed with water after 1 h, and observed immediately under a 40x oil objective of a Leica SP8 confocal microscope at magnification 0.75x (Scale = 80 μm) and magnification 3x (Scale = 20 μm). FITC fluorescence is visible as green, and chloroplast autofluorescence as magenta. (b) Magnified view of the mesophyll cells of FITC and FITC-LDH sprayed rice leaves along with line scans (white lines 1-5) and corresponding intensities of FITC fluorescence (green) and chloroplast autofluorescence (magenta).

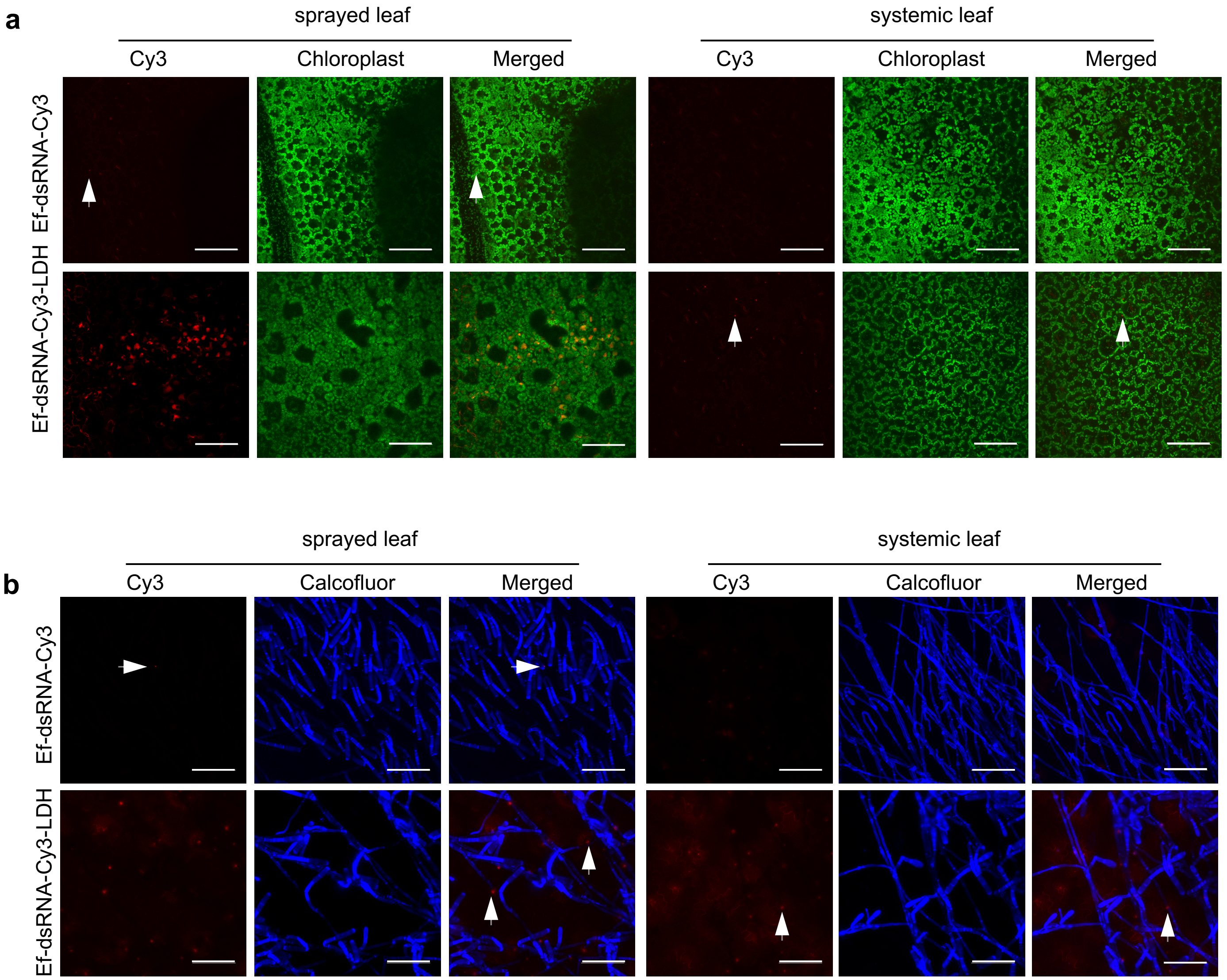


**Fig. S7 LDH facilitates systemic movement of dsRNA into newly emerged, unsprayed pea leaves.** Representative confocal images showing the presence of Cy3 signal (denoted by arrowheads) in (a) leaf cells and (b) fungal hyphae (blue) in Ef-dsRNA-Cy3 or Ef-dsRNA-Cy3-LDH sprayed (left) and unsprayed, distal (right) pea leaves. Images were captured 15 days post-spray. Cy3 fluorescence is shown in red, Calcofluor fluorescence in blue, and chloroplast autofluorescence in green; Scale = 90 μm

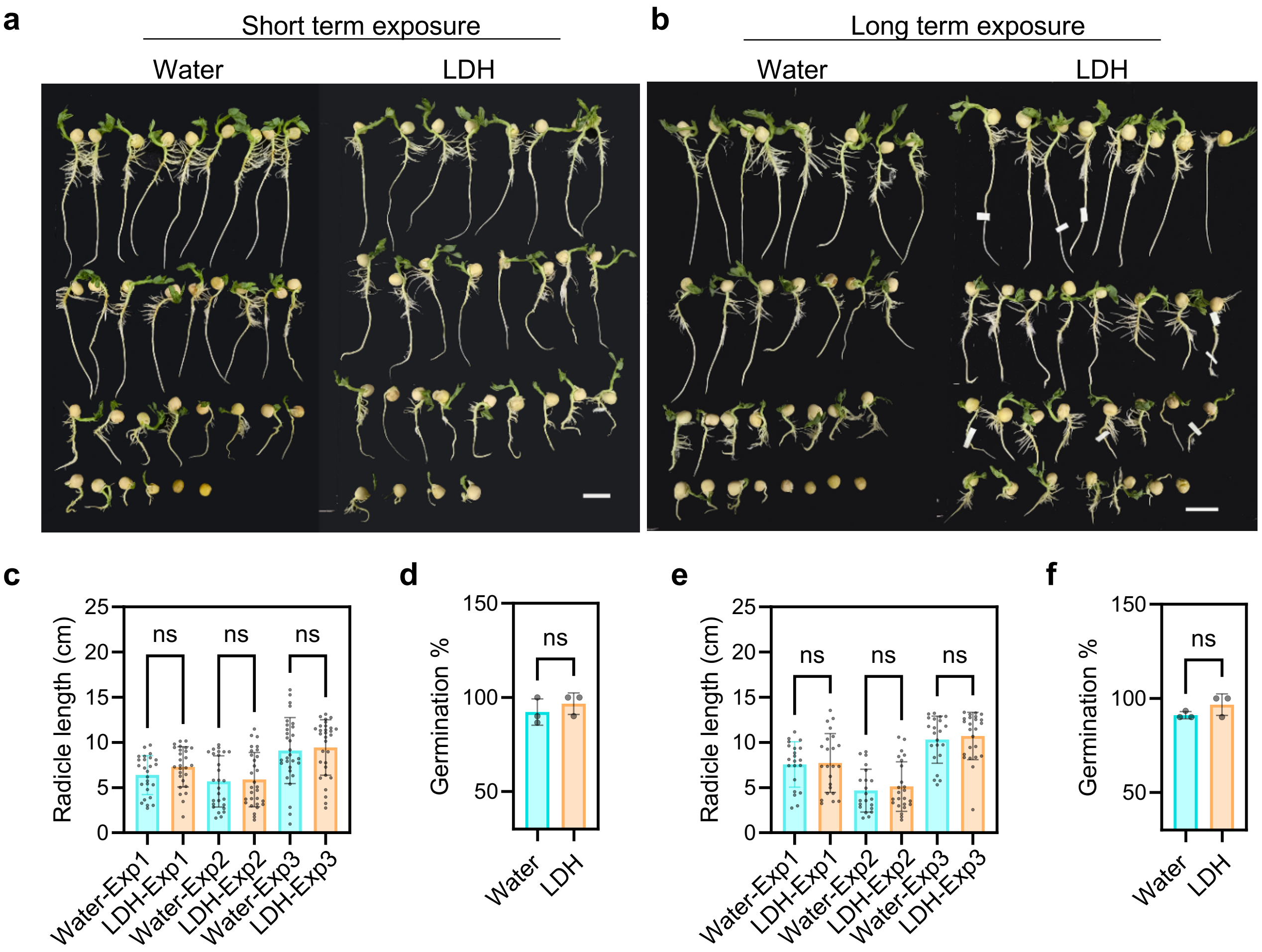


**Fig. S8 Effect of short-term and long-term exposure of MgFe-LDH on pea seed germination.** (a-b) Representative images of pea seedlings exposed to 10 mg/mL MgFe-LDH for different durations. (a) For short-term exposure, seeds were imbibed in LDH suspension for 24 h and germinated on a water-soaked cotton pad for 7 days. (b) For long-term exposure, seeds were imbibed in water for 24 h and germinated on an LDH-soaked pad for 7 days; Scale = 2 cm. (c-f) Bar graphs showing mean (±SD) radicle length and germination % of pea seeds after short-term (c-d) or long-term exposure (e-f) to LDH from three independent experiments; (n=30 seedlings per experiment). Statistical analysis was computed via One-way ANOVA ($p < 0.0001$) with Sidak's test ($\alpha < 0.05$) for radicle length and an unpaired t-test ($\alpha < 0.05$) for germination %.

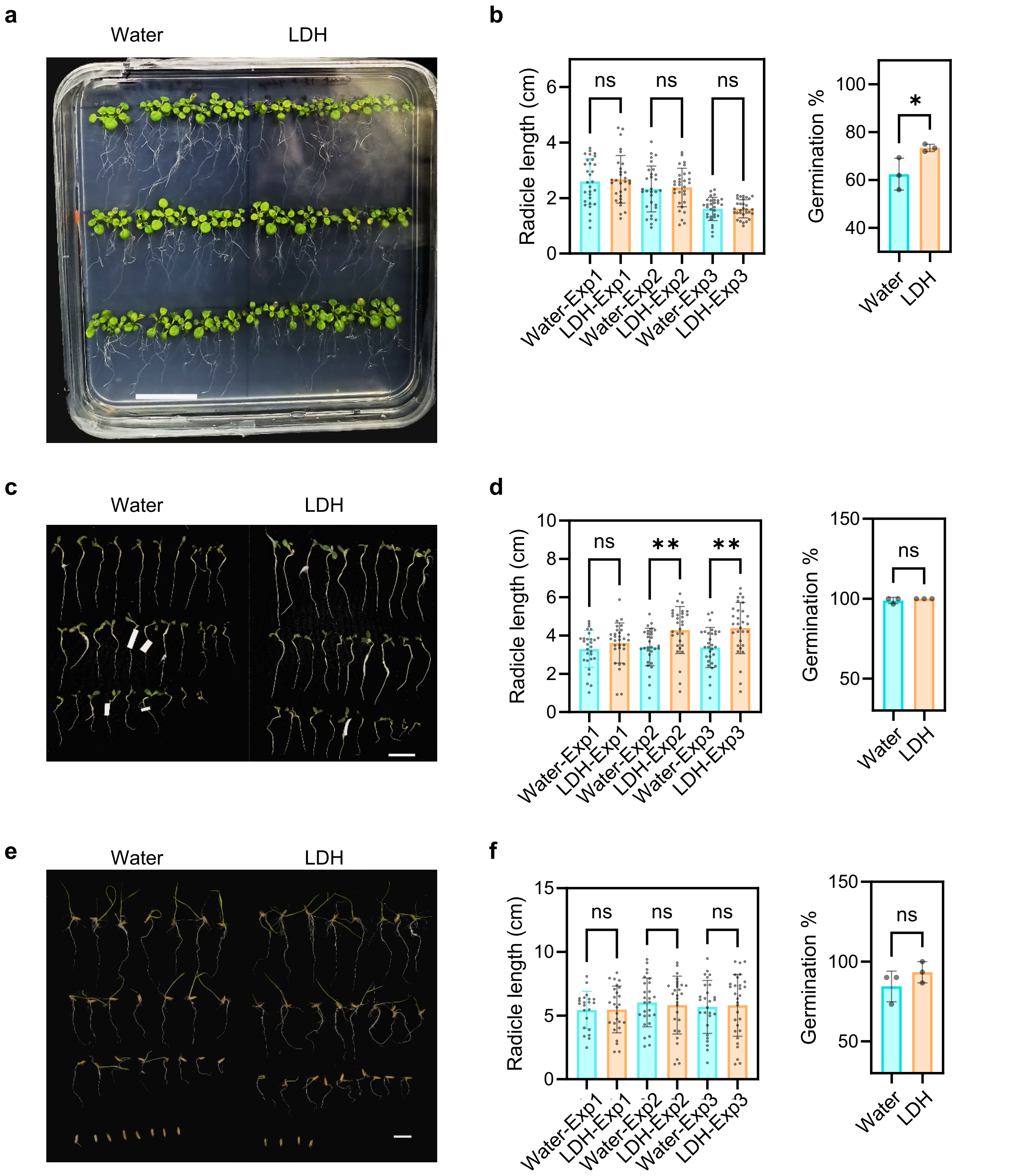


**Fig. S9 Effect of short-term exposure of MgFe-LDH on germination of Arabidopsis, *Medicago truncatula,* and rice seeds.** Representative images of (a) *Arabidopsis thaliana,* (c) *M. truncatula,* and (e) rice seedlings exposed to 4mg/mL MgFe-LDH suspension. Seeds were imbibed in the LDH suspension for 24 h and then germinated on MS media (Arabidopsis) or a water-soaked cotton pad for 7 days. n=30 per treatment; Scale = 2 cm. (b, d, f) Bar graphs showing mean (±SD) radicle length of 7 day-old-seedlings (n=30) and % germination after short-term exposure of (b) Arabidopsis, (d) *M. truncatula*, and (f) rice to LDH from three independent replicate experiments; (n=30 seedlings per experiment). Statistical analysis was computed via One-way ANOVA ($p < 0.0001$) with Sidak's test ($\alpha < 0.05$) for radicle length and an unpaired t-test ($\alpha < 0.05$) for germination %; * = $P \leq 0.05$; ** = $P \leq 0.01$, ns, non-significant

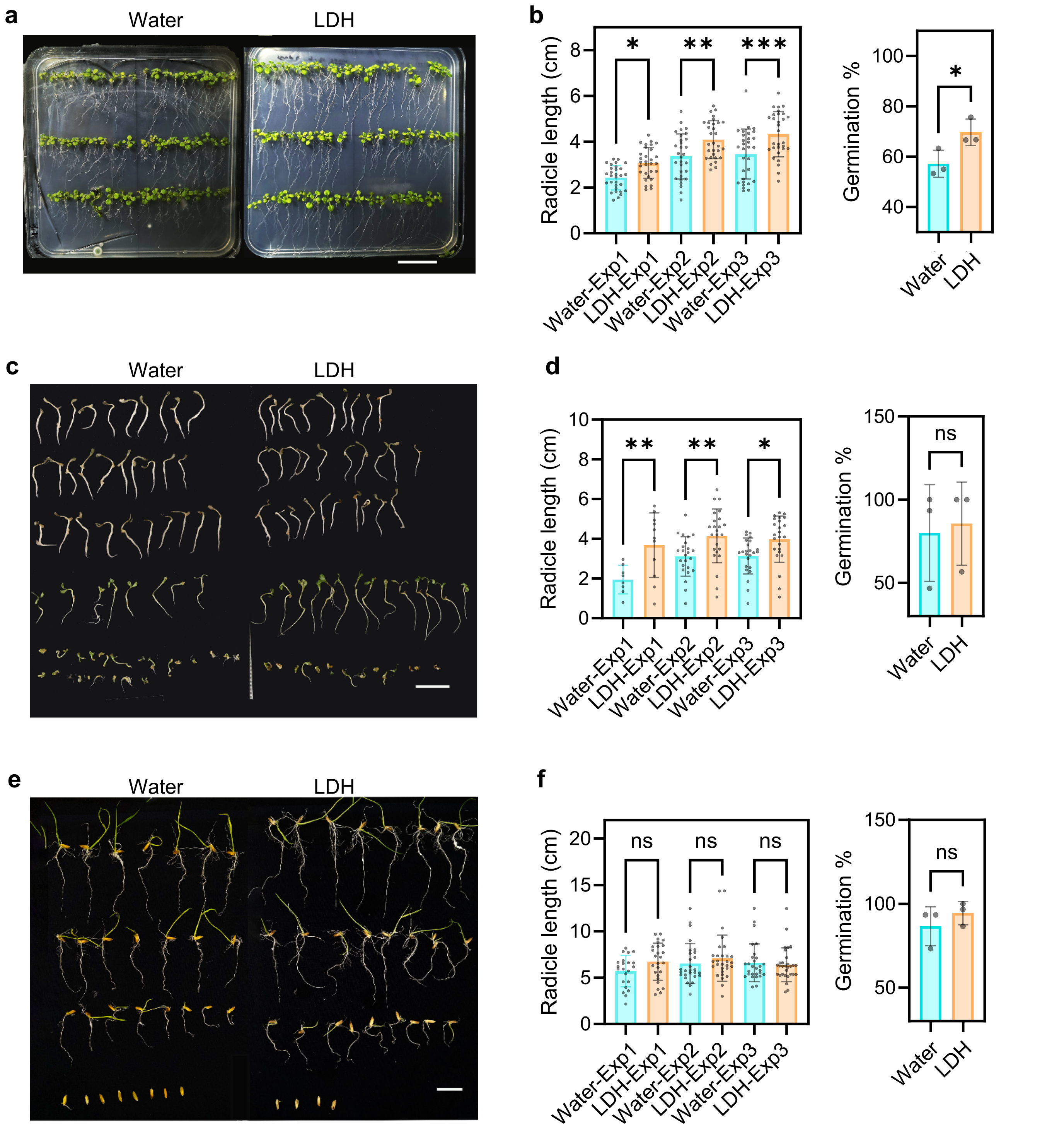


**Fig. S10 Effect of long-term exposure of MgFe-LDH on germination of Arabidopsis, *Medicago truncatula,* and rice seeds.** Representative images of (a) *Arabidopsis thaliana,* (c) *M. truncatula,* and (e) rice seedlings exposed to 10mg/mL MgFe-LDH suspension. Seeds were imbibed for 24 h in water and germinated on MS media (Arabidopsis) or a cotton pad soaked in the LDH suspension for 7 days; n=30 seedlings per treatment; Scale = 2 cm. (b, d, f) Bar graphs showing mean (±SD) radicle length of 15-day-old (Arabidopsis) or 7 day-old-seedlings (n=30) and % germination after long-term exposure of (b) Arabidopsis, (d) *M. truncatula*, and (f) rice to LDH from three independent replicate experiments; (n=30 seedlings per experiment). Statistical analysis was computed via One-way ANOVA ($p < 0.0001$) with Sidak's test ($\alpha < 0.05$) for radicle length and an unpaired t-test ($\alpha < 0.05$) for germination %. ; * = P ≤ 0.05; ** = P ≤ 0.01, *** = P ≤ 0.001; ns, non-significant.

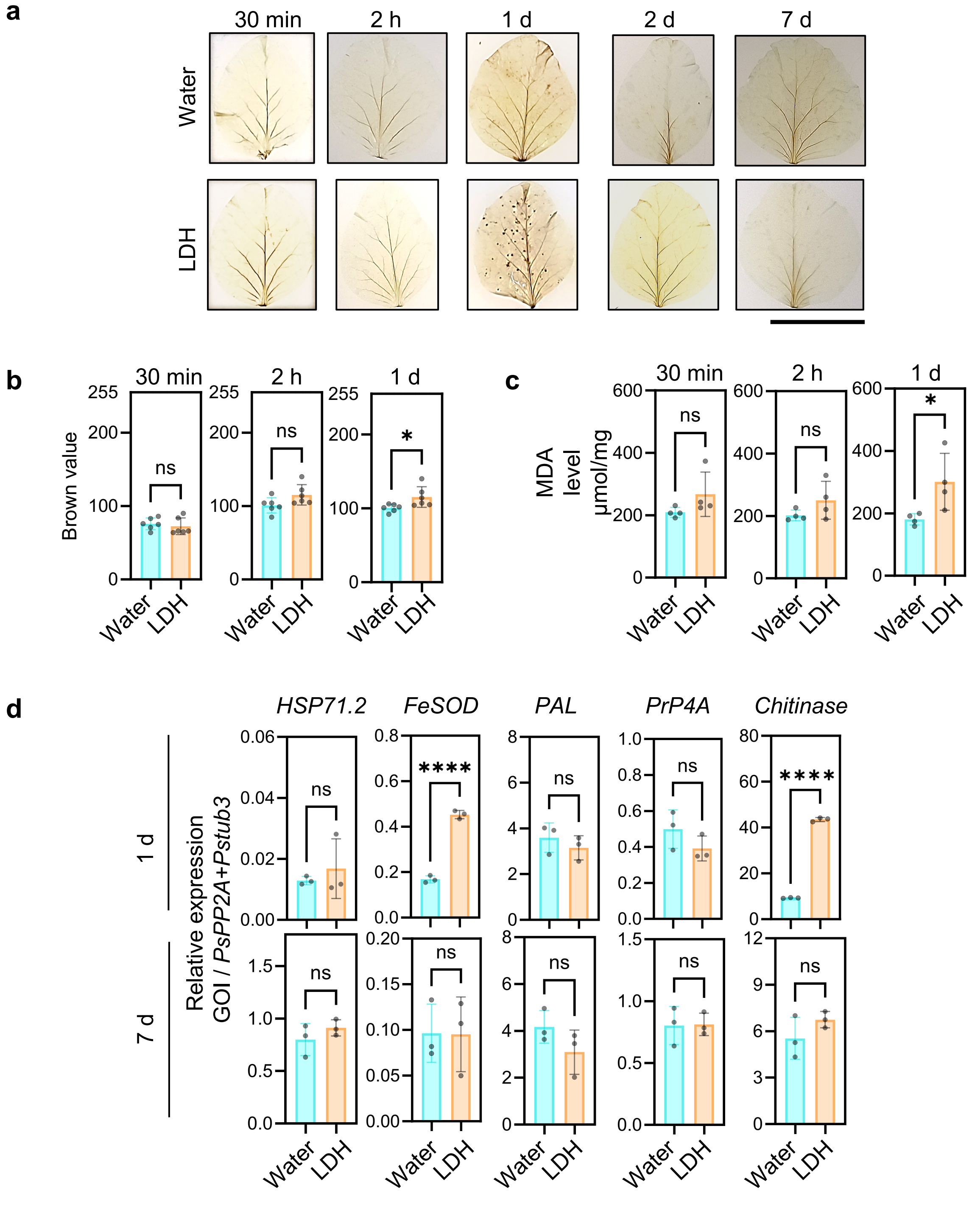


**Fig. S11 Effect of MgFe-LDH spray application on ROS generation in mature pea plants (Experiment 1).** (a) Representative DAB-stained images of pea leaves. Leaves of 20-days-old pea plants were sprayed with 10 mg/mL MgFe-LDH suspension, harvested at 30 min, 2 hours, 1 day, 2 days, and 7 days post-spray and stained with DAB to detect hydrogen peroxide accumulation; scale = 2 cm (b) Bar graphs showing mean (±SD) ROS accumulation level (Brown value 0-255) quantified using ImageJ Color-deconvolution2 plugin; n=6 leaves from three plants (c) Bar graphs showing mean (±SD) of MDA levels (μMol/mg leaf fresh weight) quantified using the thiobarbituric acid-reactive substances (TBARS) assay at 532 and 600 nm absorption; n=4 leaves from three plants. Statistical significance was computed via an unpaired t-test ($\alpha < 0.05$); ns= non-significant, * = $P \leq 0.05$. (d) Relative expression of stress marker genes normalized to the geometric mean of the endogenous controls, *Pstub3* and *PsPP2A,* at 1- and 7-days post-spray (dps) of 10 mg/mL LDH or water (n=4 pea leaves per treatment). Statistical significance was determined via unpaired t-test ($\alpha < 0.05$); **** = $P \leq 0.0001$, ns, non-significant.

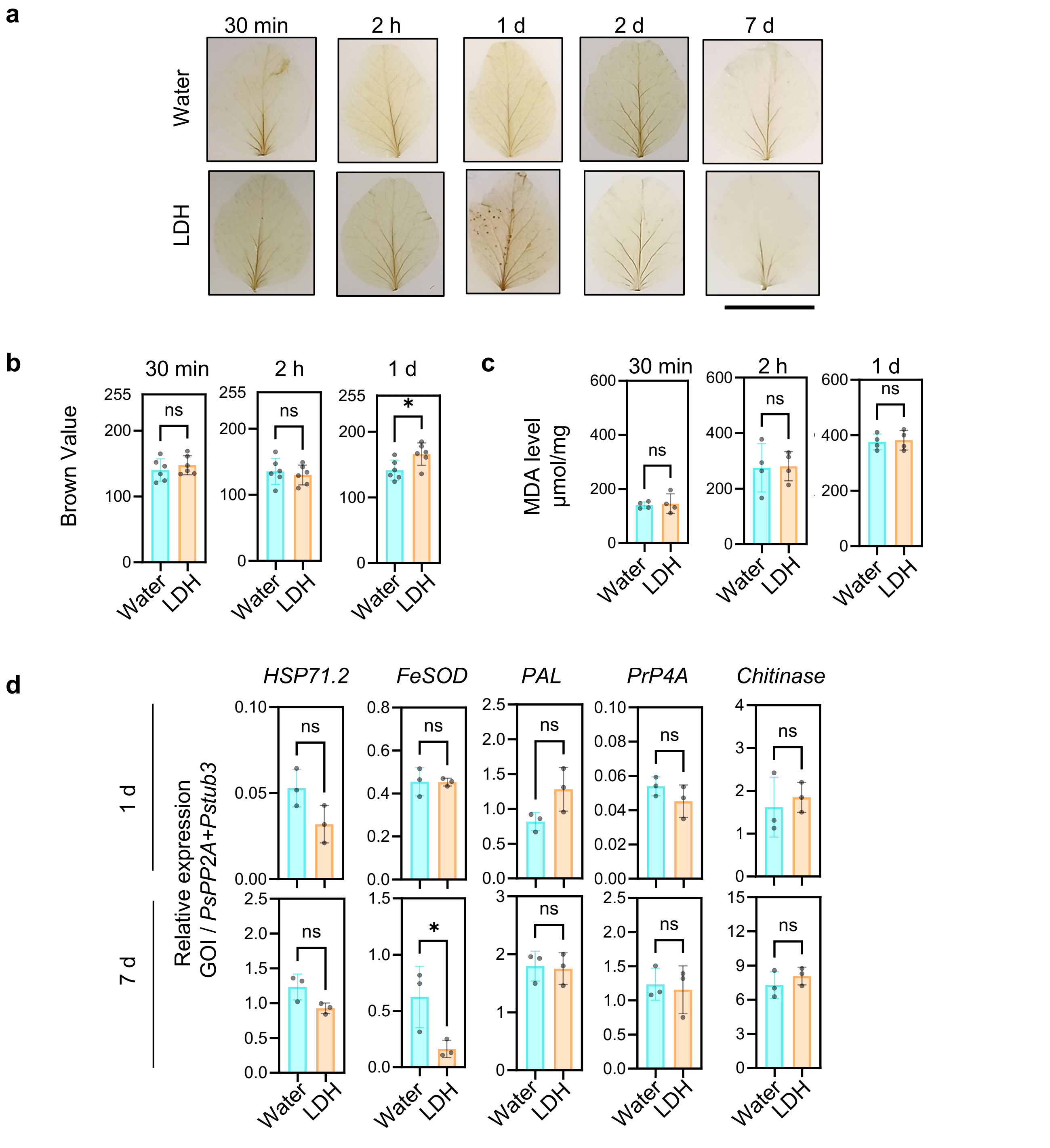


**Fig. S12 Effect of MgFe-LDH spray application on ROS generation in mature pea plants (Experiment 2).** (a) Representative DAB-stained images of pea leaves. Leaves of 20-days-old pea plants were sprayed with 10 mg/mL MgFe-LDH suspension, harvested at 30 min, 2 hours, 1 day, 2 days, and 7 days post-spray and stained with DAB to detect hydrogen peroxide accumulation; scale = 2 cm (b) Bar graphs showing mean (±SD) ROS accumulation level (Brown value 0-255) quantified using ImageJ Color-deconvolution2 plugin; n=6 leaves from three plants (c) Bar graphs showing mean (±SD) of MDA levels (μMol/mg leaf fresh weight) quantified using the thiobarbituric acid-reactive substances (TBARS) assay at 532 and 600 nm absorption; n=4 leaves from three plants. Statistical significance was computed via an unpaired t-test ($\alpha < 0.05$); ns= non-significant, * = $P \leq 0.05$. (d) Relative expression of stress marker genes normalized to the geometric mean of the endogenous controls, *Pstub3* and *PsPP2A,* at 1- and 7-days post-spray (dps) of 10 mg/mL LDH or water (n=4 pea leaves per treatment). Statistical significance was determined via unpaired t-test ($\alpha < 0.05$); * = $P \leq 0.05$, ns, non-significant.

**a**

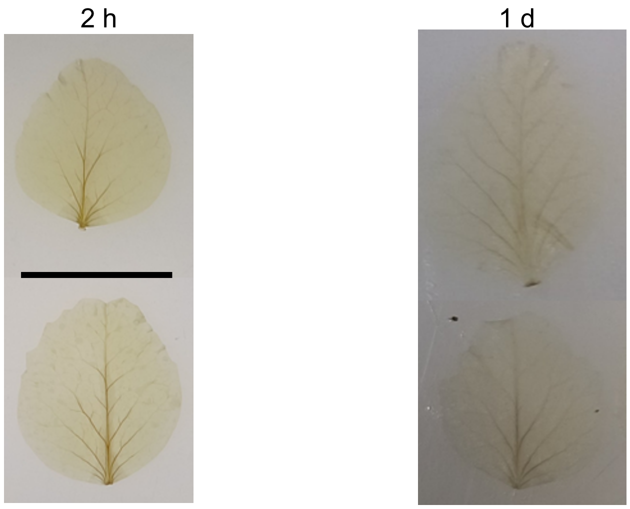


**b**

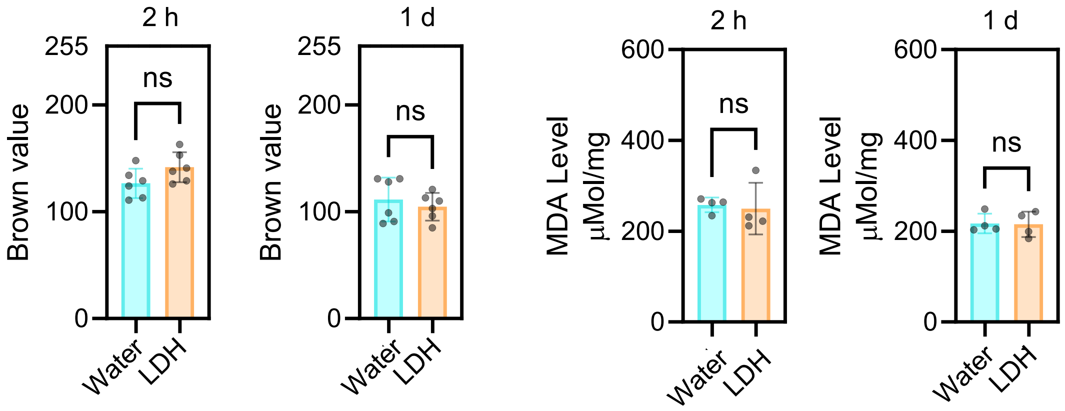


**Fig. S13 Effect of MgFe-LDH spray application on ROS generation in mature pea plants (Experiment 3).** (a) Representative DAB-stained images of pea leaves. Leaves of 20-days-old pea plants were sprayed with 10 mg/mL MgFe-LDH suspension, harvested at 2 hours and 1 day post-spray and stained with DAB to detect hydrogen peroxide accumulation; scale = 2 cm (b) Bar graphs showing mean (±SD) ROS accumulation level (Brown value 0-255) quantified using ImageJ Color-deconvolution2 plugin; n=6 leaves from three plants (c) Bar graphs showing mean (±SD) of MDA levels (µMol/mg leaf fresh weight) quantified using the thiobarbituric acid-reactive substances (TBARS) assay at 532 and 600 nm absorption; n=4 leaves from three plants.

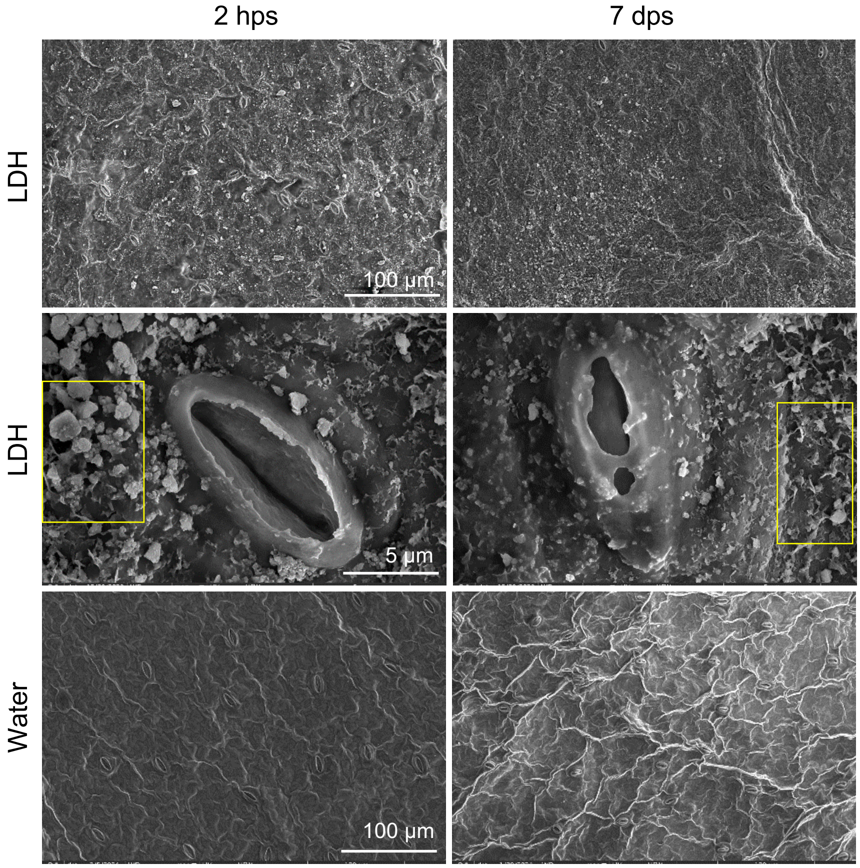


**Fig. S14 MgFe-LDH nanomaterials deposited near pea leaf stomata degrade faster than those found away from the stomata.** Representative SEM images of pea leaves sprayed with 10 mg/mL MgFe-LDH (top and middle panels) or water (bottom panel) at 2 hours post-spray (hps) and 7 days post-spray (dps). The middle panel shows a magnified view of the stomata, and the yellow boxes highlight the slower degradation of LDH nanoparticles in the regions surrounding the stomata than on the stomata.

**Methods S1**

**Characterization of the synthesized MgFe-LDH nanomaterial**

Lateral particle size was measured using Transmission electron microscopy (TEM, JEM-1400Flash), operating at 80-120 kV. High-resolution TEM (HRTEM JEM 2100 PLUS), operating at 20 kV, was used to show lattice fringes and selected area electron diffraction (SAED) patterns. Field emission scanning electron microscopy (FESEM SIGMA 500 Carl Zeiss) was used to show the surface morphology of the synthesized LDH, and the elemental composition of the MgFe-LDH was examined using colour mapping with a SEM integrated energy-dispersive X-ray spectroscopy (SEM-EDS) analyser. To determine the oxidation states of LDH, X-ray photoelectron spectroscopy (XPS) studies were conducted using an ESCALAB Xi+ instrument (Thermo Fisher Scientific), and the graphs were made in OriginPro 2024b (64-bit), 10.1.5.132. The crystal phase of the synthesized LDH in its as-prepared state was analyzed using an X-ray diffraction (XRD) PANalytical-X'pert-PRO, operated at 45 kV with a Cu-K radiation source (1.54 Å) and an X-ray diffraction angle (2θ) with a scan range of 5° to 70°. The Fourier-Transform Infrared (FTIR) spectra of the synthesised LDH were determined using a FTIR spectrophotometer (Bruker Tensor 27 FT-R) set at a range of 600–4,000 $cm^{-1}$ at a resolution of 2 $cm^{-1}$, and the sample was scanned 32 times after background correction. Brunauer–Emmett–Teller (BET) Surface analyser Quantachrome Nova-2000 was used to determine porosity and surface area utilizing the BET N2 adsorption-desorption isotherm at 77.3 K. The pore size distribution curves were examined using the B-J-H method.

**TEM analysis of dsRNA-loaded LDH**

TEM was used to visualize the morphology of the dsRNA-LDH (1:20) mixture. The mixture was sonicated for 15 min in a water bath sonicator (Athena ultrasonic bath sonicator) set at the highest power. Following sonication, the mixture was immediately suspended on a glow-discharged carbon-coated grid (Beeta Tech India, CF300-CU/50) and dried overnight at room temperature. The dried samples were imaged under a JEM-1400Flash TEM (operating at 80-120 kV).

**Effect of LDH on seed germination and mature plants**

The effect of MgFe-LDH nanoparticles on seed germination was evaluated in pea (*P. sativum*), *Arabidopsis thaliana*, *Medicago truncatula*, and rice (*Oryza sativa*). Germination percentage and radicle length were measured after treatment with water (control) or LDH. An LDH suspension of 10 mg/mL was prepared in DEPC-treated water, based on the mass loading ratio of dsRNA:LDH of 1:50 (completely loaded) for 200 µg/mL of Ef-dsRNA. Two different exposure times were tested: a) short-term exposure (24h exposure), where seeds were imbibed in water or LDH for 24 h and then germinated on a water-soaked cotton bed, and b) long-term exposure, where seeds were imbibed in water for 24 h and then placed on a water- or LDH-soaked cotton bed. The radicle length and seed germination percentages were measured 7 days post-treatment for pea, rice, and Medicago, and 15 days post-treatment for Arabidopsis. The experiment was repeated three times for each seed type.

To assess the effect of MgFe-LDH on mature plants, leaves of mature plants were sprayed with LDH suspension (10 mg $mL^{-1}$) containing 0.01% Tween-20. To determine the effect of LDH treatment on the expression of defence/stress marker genes, the relative expression of five defence/stress-related genes, including *HSP71.2* (heat shock protein 71.2)*, Chitinase, PrP4A* (pathogenesis-related protein 4A), *Fe-SOD* (Iron-superoxide dismutase), and *PAL* (phenylalanine ammonia-lyase) (Rodriguez-Serrano *et al.*, 2009), was analyzed via RT-qPCR, as described in the main Methods section. Expression values were calculated using LinRegPCR (v. 2021.2; (Ruijter *et al.*, 2009) and normalized to the reference genes *Ps Protein Phosphatase 2A* (*PsPP2A*; NM_001427333.1) and *Pstubulin*. Primer sequences are listed in **Table S1**.

To determine the effect of LDH treatment on hydrogen peroxide accumulation, control and LDH-treated leaves were harvested at 30 min, 2 h, and 1 day post-spray and stained with 1mg $mL^{-1}$ acidified DAB (3,3′-diaminobenzidine, Himedia) as per the protocols described in Thordal-Christensen *et al.* (2002) and Daudi & O'Brien (2012). DAB staining intensity was quantified using the ImageJ (version 1.54f) color_deconvolution2 plugin (https://github.com/landinig/IJ-Colour_Deconvolution2/blob/main/colour_deconvolution2.jar) and selecting the H-DAB to isolate DAB staining. The leaf area was marked using the ImageJ freehand tool, and the mean intensity of brown value, ranging between 0 and 255 pixel intensity, was measured. Here, 0 represents darkest brown, and 255 represents white, with the intermediate values reflecting a gradient of brown colour from minimum to maximum intensity. The resulting brown values for control and LDH-treated samples were recorded for each leaf after background correction. Each value was then subtracted from 255 to invert the range such that a higher value represents a higher DAB staining intensity. The values were plotted as mean (±SD) in GraphPad Prism (v10.3.0).

To determine the effect of LDH treatment on Malondialdehyde (MDA) accumulation, MDA was quantified in control and LDH-treated leaves harvested at 30 min, 2 h, and 1 day post-spray, as described in (Heath & Packer, 1968). Briefly, harvested pea leaves were homogenized with liquid nitrogen, followed by the addition of 0.1% Trichloroacetic acid (TCA). The mixture was centrifuged at 15000 x g, at 4°C for 10 min, and the supernatant was collected and diluted with 0.5 % Thiobarbituric Acid (TBA). The solution was heated at 95 °C for 25 min in a water bath, followed by immediate cooling on ice. The resulting Thio-barbituric acid-reactive substances (TBARS) were quantified by measuring absorbance at 532 and 600 nm with a spectrophotometer. The absorbance values at 532 nm were subtracted from those recorded at 600 nm, and the MDA concentration was calculated using the Lambert-Beer law with an extinction coefficient εM= 155 $mM^{-1}cm^{-1}$. Results are presented as μmols MDA $g^{-1}$ FW and plotted using GraphPad Prism (v10.3.0).

**Identification of optimal dsRNA dose for foliar spray application**

Ef-dsRNA was prepared at 200 and 500 ng/μL concentrations. Each dose was mixed with Tween 20 (Sigma-Aldrich, Cat #9005-64-5) at a final concentration of 0.01% and sprayed onto individual leaves of pea plants (*Pisum sativum*) using an airbrush. After 24 h, the same leaves were spray inoculated with 10,000 conidia/mL, as previously described (Ray & Chandran, 2024). Leaves were harvested at 3-, 5-, and 7-days post-inoculation (dpi). At 3 dpi, the number of primary and secondary fungal hyphae was quantified after staining fungal colonies with trypan blue. At 7 dpi, the percentage of each leaf covered by powdery mildew symptoms was measured using ImageJ (version 1.54f) software with a dedicated plugin for fungal structure analysis (Ray & Chandran, 2024). The relative expression levels of *Ef* and *Epβ-tubulin* were quantified at 3, 5, and 7 days post-inoculation (dpi) using RT-qPCR and normalized to the internal control genes, *Pisum sativum Protein Phosphatase 2A* (*PP2A*; NM_001427333.1) and *PsTubulin3*.

**Characterization of FITC-labelled MgFe-LDH via DLS, TEM and FTIR**

For the DLS assay, 10 μL of the FITC-LDH (1:20) suspension was diluted 1000x and probe sonicated (Labman PRO656 - Probe Sonicator-Touch screen) for 2 min. 800 μL of FITC-LDH or LDH suspension was added to a Folded Capillary cell (DTS1070, Malvern Zetasizer) for size and surface charge analysis in a DLS machine (Malvern Nano Zetasizer) at 25°C and 90° angle. The assay was repeated with three independent batches of FITC-LDH, and the mean values (±SD) were plotted in OriginPro 2024b (64-bit), v10.1.5.132. The mean (±SD) nanomaterial hydrodynamic size (nm) was plotted on the left Y-axis, and the mean (±SD) Zeta or surface charge (mV) on the nanomaterial was plotted on the right Y-axis. The sample was also analysed via TEM microscopy on a glow-discharged carbon-coated grid (Beeta Tech India, CF300-CU/50).

For the FTIR assay, the FITC-LDH suspension was dried at 37 °C for 3 days until all the water had completely evaporated. The dried powder was homogenized with a micro pestle and analysed in an FTIR Spectrometer (Bruker Tensor 27 FT-IR). The instrument was set to a range of 600–4,000 $cm^{-1}$ at a resolution of 2 $cm^{-1}$, and the sample was scanned 32 times after background correction. The transmittance % (Y-axis) was then plotted as a linear graph in OriginPro 2024b (64-bit), v10.1.5.132 by subtracting the baseline and plotting the wavenumber ($cm^{-1}$) on the X-axis. The characteristic wavenumbers are shown above the vertical dotted lines.

**Uptake of FITC-LDH in *Nicotiana benthamiana* and rice leaves**

FITC-LDH (1:20) and FITC (10 µg/mL) suspensions were sprayed on leaves of *Nicotiana benthamiana* (5-7 weeks old) and rice (10-12 weeks old) plants. Following 1 h incubation in the dark, the FITC-LDH and FITC-sprayed leaves were washed with autoclaved Milli-Q water and blot-dried with Kim wipes. The FITC and FITC-LDH-sprayed Nicotiana leaves (1 h post-spray) were sectioned into 0.5 $cm^2$ square pieces and visualized under the 63x oil objective of a Leica SP8 CLSM. Images were captured at two magnifications: 0.75x and 2x. For rice, 0.5 $cm^2$ square leaf sections (1 h post spray) were visualized under the 40x oil objective of a Leica SP8 CLSM, and images were captured at two magnifications: 0.75x and 3x. The laser settings and image acquisition and analysis protocols used for pea (described in the main methods section) were also used here.

**Systemic movement of Ef-dsRNA into unsprayed distal leaves**

To visualize dsRNA-Cy3 signals in sprayed and newly emerged unsprayed upper leaves at 15 days post-spray, leaf sections were taken from uninfected plants, and imaged using a Leica SP8 confocal laser scanning microscope (CLSM) as previously described. To visualize dsRNA-Cy3 signals within fungal structures at 15 days post-spray or 14 days post-inoculation (dpi), sections of infected sprayed and newly emerged unsprayed leaves were stained with Calcofluor White and imaged on a Leica SP8 CLSM. Z-stacks (0.35 µm thickness) were captured and processed using the Leica Application Suite X.

**References**


DAUDI, A. & O'BRIEN, J. A. 2012. Detection of hydrogen peroxide by DAB staining in Arabidopsis leaves. *Bio Protoc,* 2**,** e263.

HEATH, R. L. & PACKER, L. 1968. Photoperoxidation in Isolated Chloroplasts: I. Kinetics and Stoichiometry of Fatty Acid Peroxidation. *Arch Biochem Biophys,* 125**,** 189-198.

RAY, P. & CHANDRAN, D. 2024. Spray inoculation and image analysis-based quantification of powdery mildew disease severity on pea leaves. *MethodsX,* 13**,** 102980.

RODRIGUEZ-SERRANO, M., ROMERO-PUERTAS, M. C., PAZMINO, D. M., TESTILLANO, P. S., RISUENO, M. C., DEL RIO, L. A. & SANDALIO, L. M. 2009. Cellular response of pea plants to cadmium toxicity: cross talk between reactive oxygen species, nitric oxide, and calcium. *Plant Physiol,* 150**,** 229-43.

RUIJTER, J. M., RAMAKERS, C., HOOGAARS, W. M., KARLEN, Y., BAKKER, O., VAN DEN HOFF, M. J. & MOORMAN, A. F. 2009. Amplification efficiency: linking baseline and bias in the analysis of quantitative PCR data. *Nucleic Acids Res,* 37, e45.

THORDAL-CHRISTENSEN, H., ZHANG, Z., WEI, Y. & COLLINGE, D. B. 2002. Subcellular localization of $H_2O_2$ in plants. $H_2O_2$ accumulation in papillae and hypersensitive response during the barley—powdery mildew interaction. *Plant J,* 11, 1187-1194.
